# Structurally Constrained Functional Connectivity Reveals Efficient Visuomotor Decision-Making Mechanisms in Action Video Gamers

**DOI:** 10.1101/2025.06.13.659455

**Authors:** Kyle Cahill, Mukesh Dhamala

**Affiliations:** Department of Physics and Astronomy, Georgia State University, Atlanta, GA, USA; Neuroscience Institute, Georgia State University, Atlanta, GA, USA; Center for Behavioral Neuroscience, Center for Diagnostics and Therapeutics, Georgia State University, Atlanta, GA, USA; Tri-Institutional Center for Translational Research in Neuroimaging and Data Science (TReNDS), Georgia State University, Georgia Institute of Technology, and Emory University, Atlanta, GA, USA

## Abstract

Long-term action video game (AVG) playing has been linked to improved response times (∼190 ms) without accuracy tradeoffs in time-sensitive visuomotor decisions, but how it reshapes neural circuits that enable this behavioral advantage is unclear. In this study, Cognitive Resource Reallocation (CRR) is introduced as a candidate mechanism for how sustained engagement with AVGs drives behaviorally relevant neuroplasticity through neuroplastic refinement. Using the AAL3 structural connectivity atlas, we apply structural constraints to functional connectivity (SC-FC) and directed functional connectivity (SC-dFC) in gamers and non-gamers. Our results provide strong support for the CRR hypothesis and demonstrate that the brain plausibly reallocates cognitive resources over time to optimize task-relevant networks in high-demand environments such as AVGs, enhancing the integration of contextual information and refining motion processing, which may be a key mechanism in explaining more efficient visuomotor decision-making. These findings position action video games as powerful tools for studying experience-driven neuroplasticity, with implications for cognitive training, rehabilitation, and optimizing real-world visuomotor decisions.

## 1. Introduction

Video games have become a ubiquitous form of entertainment worldwide, with over 3.2 billion gamers globally in 2023 ^1^, and the video game industry has continued to expand at a rapid pace ^2, 3^. This cultural shift is not only transforming leisure activities but is also driving a growing body of research into the cognitive and neural impacts of video game play. Action-based video games, such as First-Person Shooters (FPS) and Real-Time Strategy (RTS) games, have garnered attention for their potential cognitive benefits, with studies reporting a broad range of improvements across various cognitive domains.^4–10^ These include enhanced sensorimotor integration^11^, attentional control,^12–14^ executive function^15^, cognitive flexibility^16, 17^ in regular players.

These cognitive enhancements extend beyond video gaming and have been shown to transfer to real-world applications, including surgery,^18^ driving^15^, military training^19^, and aviation^20, 21^. In addition to these observed cognitive benefits, action video games are increasingly being explored as clinical tools, with the FDA-approved EndeavorRx improving attention and self-regulation in children with ADHD.^22^ Action video games are also being shown to enhance higher-order cognitive functions, such as attentional control and reading performance, even in individuals with neurodevelopmental conditions like dyslexia, underscoring their potential for cognitive training and rehabilitation.^23^

Beyond cognitive benefits and rehabilitation potential, an increasing number of neuroimaging studies have demonstrated structural and functional adaptations associated with action video gameplay^10, 11, 24–33^ These studies have reported changes in gray matter volume,^30, 34^ cortical thickness^35^, white matter structure^33, 36, 37^, and large-scale functional connectivity patterns^10, 28, 31^ particularly in networks supporting visuospatial cognition^38^, and attention.^26, 28^ However, only a few studies have integrated both structural and functional MRI data within the same analysis, less so in healthy, non-addicted gamers^10, 26, 31^ Moreover, while neuroimaging evidence supports widespread plasticity in gamers, many of these studies have lacked behavioral validation via direct cognitive assessments, making it unclear as to which connectivity differences translate into measurable cognitive advantages.^31^ As a result, the relationship between specific neural adaptations and behavioral performance remains an open question, particularly in how functional network coordination operates within the fundamental constraint that rapid interregional communication is facilitated by white matter tracts.^39^

The mechanisms of neuroplasticity, reflecting the brain’s ability to reorganize its structure and function, have been extensively studied at the mesoscale^40^ encompassing processes such as Hebbian plasticity, long-term potentiation (LTP), synaptic pruning, and homeostatic plasticity, alongside broader mechanisms like neurogenesis and myelination, a unified framework by which these mechanisms coordinate to induce large-scale effects remains elusive.

This study introduces Cognitive Resource Reallocation (CRR) as a potentially unifying framework to explain how mesoscale neuroplastic processes collectively give rise to large-scale effects in action video game players. The CRR hypothesis posits that the brain dynamically reallocates cognitive resources to anatomically plausible, functionally relevant regions and pathways in response to task demands and environmental pressures. Rather than specifying the exact mechanistic contributions of individual mesoscale neuroplastic processes, CRR provides an organizing principle for how these mesoscale changes may be organized to produce large-scale neuroplastic adaptations. This study tests whether observed connectivity shifts align with CRR’s predictions, offering a framework for evaluating its role in experience-driven neuroplasticity.

In the context of action video games, CRR would progressively optimize neural mechanisms along anatomically viable pathways that support proficient gameplay in fast-paced, sensory-rich, high-pressure environments. A key example is visuomotor decision-making, where failure to make swift and accurate decisions in response to salient visual cues leads to errors that hinder performance. Over time, sustained engagement in action video games enforces a baseline level of visuomotor efficiency, which is essential for maintaining moderate gameplay success. As a result, cognitive functions that support efficient visuomotor decision-making, such as visual processing, visuomotor integration, attentional control, and cognitive flexibility, along with their underlying neural substrates, are expected to improve as a byproduct of prolonged gameplay. Understanding how the brain facilitates these neural adaptations in response to long-term action video game experiences is crucial for identifying the neurobiological mechanisms that drive cognitive enhancements.

Our previous findings have demonstrated that regular long-term action video game players exhibit more efficient visuomotor decision-making, demonstrating ∼190ms faster response times without sacrificing accuracy.^32^ This behavioral advantage is clear, and while these changes are expected to occur along anatomically plausible, functionally relevant pathways, a comprehensive understanding of the neural mechanisms driving these optimizations remains unknown.

To test CRR, we constrain both undirected and directed functional connectivity in action gamers and non-gamers by applying anatomical constraints from validated white matter tracts between brain regions. This allows us to construct structurally constrained undirected (SC-FC) and directed (SC-dFC) connectivity matrices. While functional connectivity alone provides valuable insights, integrating it with structural connectivity data ensures that the observed functional interactions are constrained by biologically plausible anatomical pathways, enhancing both reliability and interpretability.^41^ This approach allows us to investigate how long-term video game playing influences brain connectivity in anatomically feasible connections, while also assessing their behavioral relevance and their global and local network properties. Additionally, we incorporate SC-dFC, assessed via Granger causality, to examine directed interactions between regions. By considering both SC-FC (undirected synchrony) and SC-dFC (directed causal interactions), we ensure that our analysis captures both functionally plausible neural synchrony and causal information flow, all within the constraints of underlying structural connectivity.^42, 43^ This approach provides a structurally informed measure of effective connectivity, offering a more complete picture of how long-term action video game experience shapes neural communication along anatomically feasible white matter pathways.

This study advances our understanding of experience-induced neuroplasticity by investigating how long-term action video game play enhances visuomotor decision-making efficiency without sacrificing accuracy.^32^ By testing CRR as a clear and falsifiable theoretical framework, we aim to assess whether it is a robust explanatory principle for behaviorally induced neural adaptations. Our findings may further elucidate the mechanisms, by which video games function as cognitive training tools and strengthen their validity as a rigorous experimental medium for studying experience-dependent neuroplasticity.

## 2. Results

### 2.1 Structurally Constrained Whole-Brain Functional and Directed Connectivity Differences

To investigate large-scale task-relevant connectivity adaptations associated with action video game playing, we examined structurally constrained functional connectivity (SC-FC) and structurally constrained directed functional connectivity (SC-dFC) between long-term action video gamers and non-gamers. By constraining our analysis with underlying white matter pathways, this approach ensures that observed group differences reflect meaningful adaptations rather than arbitrary or spurious connections.

The SC-FC results of the significant connections are presented as a heat map of mean differences between groups, displayed as a connectivity matrix in Figure 1a. Connections where gamers showed stronger structurally constrained functional connectivity are indicated by warm colors (red, orange), whereas connections stronger in non-gamers are indicated by cool colors (cyan, blue). Gamers exhibited a significantly greater number of enhanced (*p* < 0.05) SC-FC connections compared to non-gamers (278 ± 17 vs. 220 ± 15; *Z* = 2.60, *p* < 0.01). Significance was determined by a Gaussian approximation to estimate the standard error of total connection counts. SC-FC analyses revealed greater connectivity in gamers across occipital-limbic, occipital-parietal, frontal-limbic, and frontal-parietal pathways. In contrast, non-gamers exhibited stronger SC-FC between frontal-occipital regions and within the cerebellum (Figure 1a).

**Figure 1.**
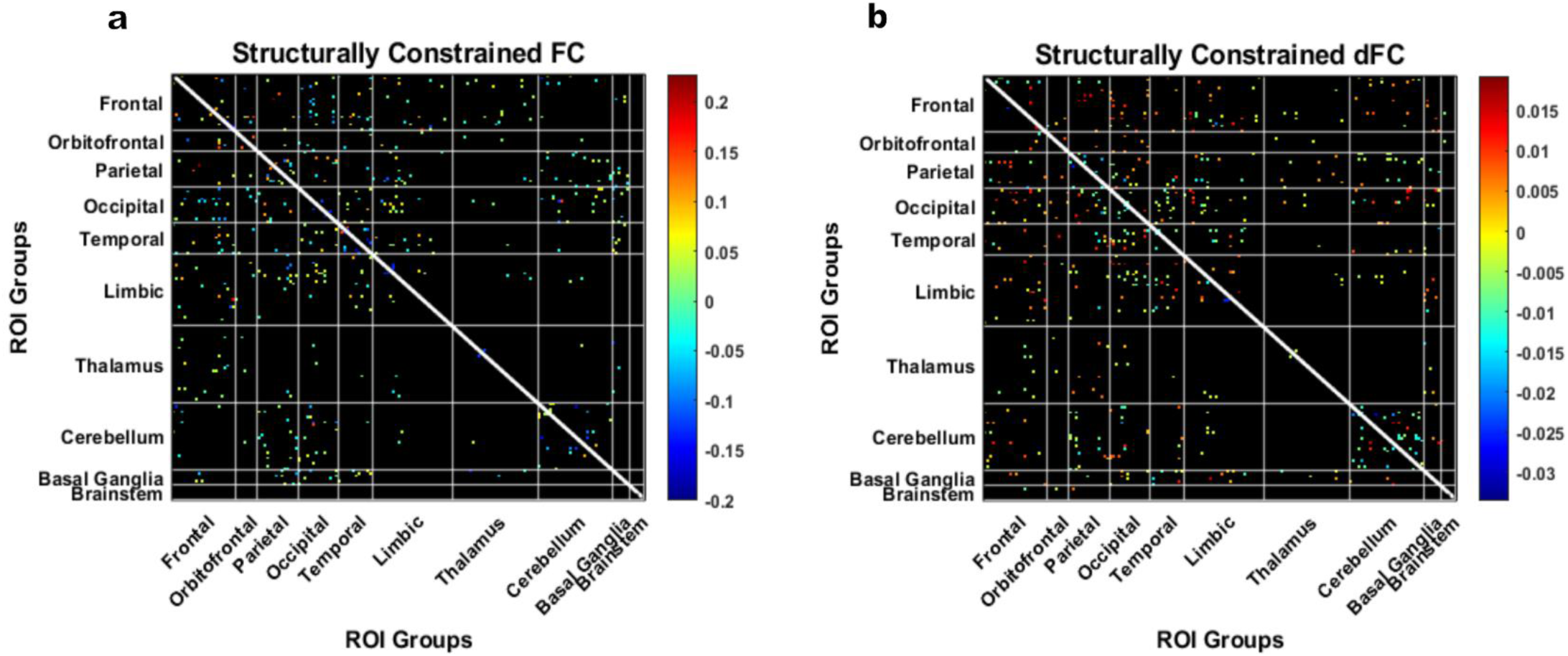
Structurally Constrained Functional Connectivity & Directed Connectivity Mean Differences Between Gamers and Non-Gamers. (a) SC-FC (-) group differences, where positive values indicate greater connectivity in gamers and negative values indicate stronger connectivity in non-gamers (*p* < 0.05). (b) SC-dFC (→) group differences measured using time-domain Granger causality (TGC) to capture effective connectivity (*p* < 0.05).

The SC-dFC results are shown in Figure 1b as a heat map of mean group differences, using the same matrix format as Figure 1a. Although non-gamers exhibited a greater total number of significantly stronger SC-dFC (313 ± 18 vs. 249 ± 16; *Z* = 2.70, *p* < 0.01), gamers showed greater SC-dFC between frontal and occipital regions, suggestive of more targeted top-down visual processing. Non-gamers showed significantly greater SC-dFC within cerebellar regions, consistent with the SC-FC findings and reinforcing a distinct inter-cerebellar profile. A detailed breakdown of the AAL3 atlas regions, categorized by functional divisions, is available in Supplementary Figure 1.

### 2.2 Brain-Behavior Relationships Between Connectivity and Response Times

To assess the behavioral relevance of SC-FC and SC-dFC connections, we examined correlations between SC-FC, SC-dFC, and response times (RTs) across both groups (see Figure 2).

**Figure 2.**
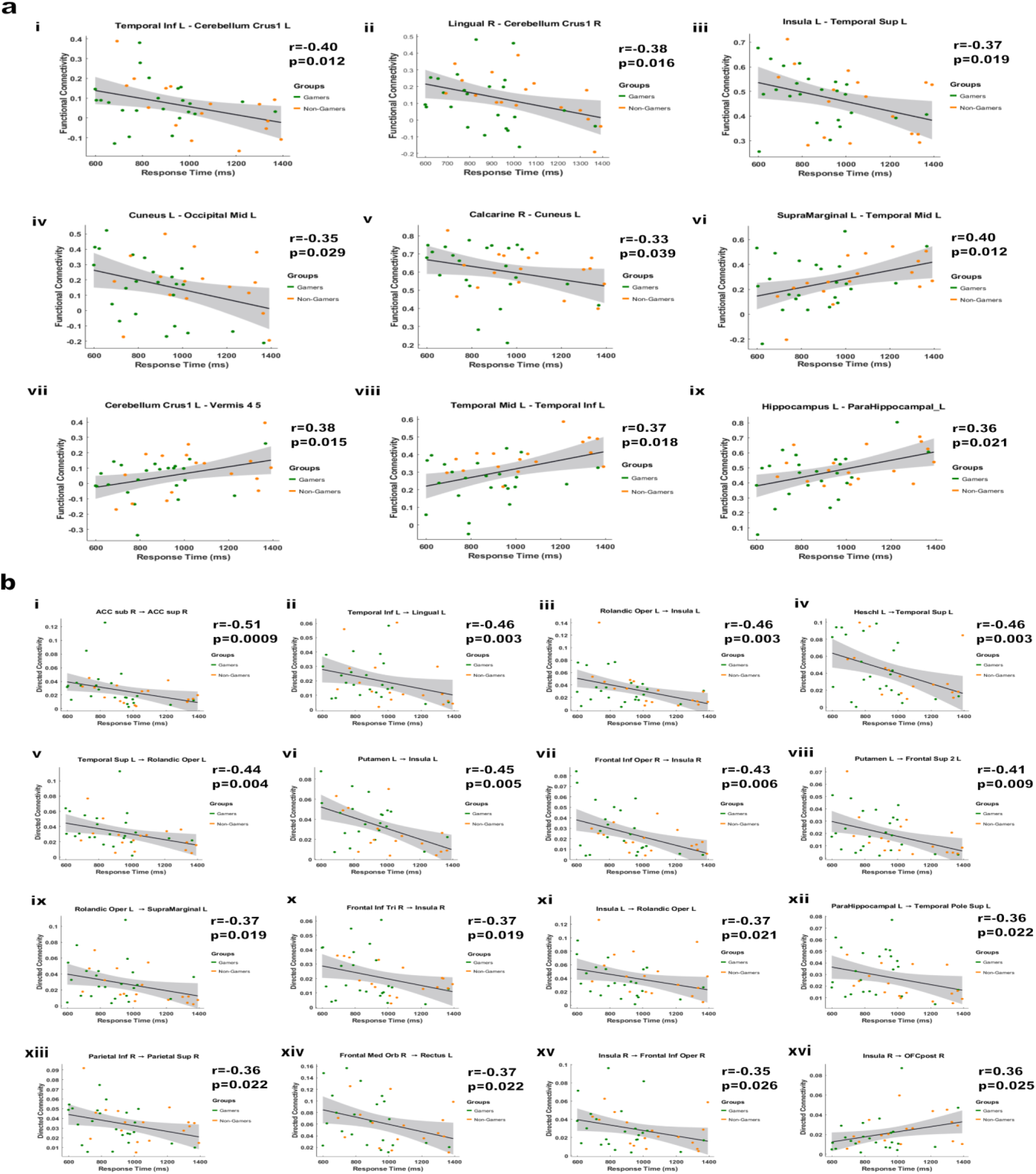
Brain-Behavior Relationships Between Functional Connectivity Measures and Response Times. (a) Significant correlations between structurally constrained functional connectivity (SC-FC) and response times, ranked from lowest to highest p-values, separated by (i-v) negative and (vi-ix) positive correlation coefficients. Negative correlations reflect connections where increased SC-FC predicts faster decision-making, while positive correlations indicate connections where stronger SC-FC is associated with slower response times. (b) Significant correlations between structurally constrained directed functional connectivity (SC-dFC) and response times, ranked from lowest to highest p-values, separated by negative (i-xv) and positive (xiv) correlation coefficients, capturing effective connectivity linked to task performance (*p < 0.05*).

#### 2.2.1 SC-FC (-) Correlations with Response Times

Several SC-FC pairwise relationships showed significant negative correlations with RT, indicating that stronger connectivity was associated with faster performance. These included connections between the left inferior temporal and left cerebellum crus I (*r* = –0.40, *p* = 0.012), right lingual and right cerebellum crus I (*r* = –0.38, *p* = 0.016), and left insula and left superior temporal cortex (*r* = –0.37, *p* = 0.019). Additional correlations were observed in early visual areas, including left cuneus – left middle occipital (*r* = –0.35, *p* = 0.029) and right calcarine – left cuneus (*r* = –0.33, *p* = 0.039). The cuneus, positioned just above the calcarine sulcus, is thought to play a key role in routing visual input into the dorsal stream^44, 45^. The cuneus’ involvement here suggests that faster responders may engage more early-stage dorsal relays for visuomotor integration.

There were also several SC-FC relationships that showed significant positive correlations with RT, indicating that stronger connectivity was associated with slower performance. These included left supramarginal – left middle temporal (*r* = 0.40, *p* = 0.012), left cerebellum crus I – Vermis 4,5 (*r* = 0.38, *p* = 0.015), and left middle temporal – left inferior temporal (*r* = 0.37, *p* = 0.018). Additionally, SC-FC between the left hippocampus and left parahippocampus was positively correlated with response time (*r* = 0.36, *p* = 0.021).

#### 2.2.2 SC-dFC (→) Correlations with Response Times

A wide array of effective pairwise causal relationships given by SC-dFC was negatively correlated with RT. As shown in Figure 2b, the strongest correlation was observed between the right subgenual and supracallosal anterior cingulate cortex (r = –0.51, *p* = 0.0009). Additional SC-dFC relationships associated with faster response times included left temporal middle → left lingual (*r* = –0.46, *p* = 0.003), left rolandic operculum → left insula (*r* = –0.46, *p* = 0.003), left insula → left superior temporal (*r* = –0.46, *p* = 0.003), and left superior temporal → left rolandic operculum (*r* = –0.44, *p* = 0.004).

Several subcortical and frontal pathways were also significant, including left putamen → left insula (*r* = –0.45, *p* = 0.005), left frontal operculum → left insula (*r* = –0.43, *p* = 0.006), and left putamen → left superior frontal gyrus (*r* = –0.41, *p* = 0.009). Right hemisphere relationships included right frontal inferior triangularis → right insula (*r* = –0.37, *p* = 0.019), right frontal inferior orbitalis → right rolandic operculum (*r* = –0.37, *p* = 0.021), and right insula → right frontal inferior operculum (*r* = –0.35, *p* = 0.026).

Additional significant SC-dFC findings included right rolandic operculum → left supramarginal (*r* = –0.37, *p* = 0.019), left superior parietal → itself (*r* = –0.36, *p* = 0.022), right frontal medial orbital → right rectus (*r* = – 0.37, *p* = 0.022), and left parahippocampus → left superior temporal pole (*r* = –0.36, *p* = 0.022). One relationship, right insula → right posterior orbitofrontal cortex (*r* = 0.36, *p* = 0.025), showed a significant positive correlation with RT.

### 2.3 Behaviorally Relevant Group Differences in SC-FC and SC-dFC

Among the behaviorally relevant SC-FC connections, a significant group difference was observed between the left middle temporal and inferior temporal gyri. This connection was significantly stronger in non-gamers (*p* = 0.002) as shown in Figure 3a(i) and showed a positive correlation with response times (*r* = 0.37, *p* = 0.018), which is displayed in Figure 2a(vii).

**Figure 3.**
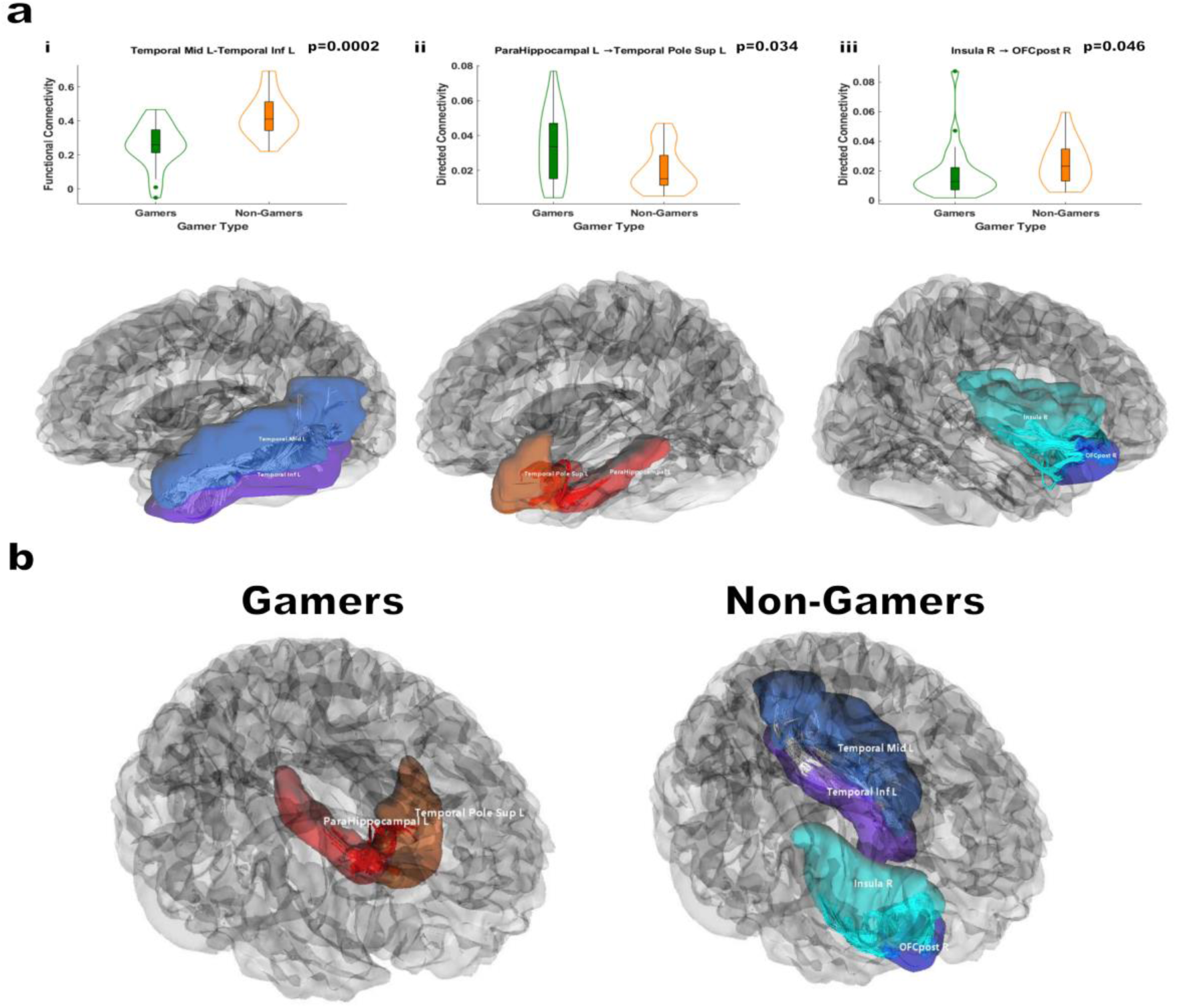
Behaviorally Relevant Connectivity Differences Between Gamers and Non-Gamers. (a) Violin plots comparing functional (i) and directed (ii, iii) connectivity for key brain regions, including 211 Temporal Mid L – Temporal Inf L, Parahippocampal L →Temporal Pole Sup L, and Insula R →OFCpost R, with significant group differences indicated by p-values. (b) 3D renderings of the respective regions for Gamers (left) and Non-Gamers (right), highlighting the anatomical locations where significant connectivity differences were observed. The brain regions shown are left mid-temporal, left inferior temporal, left parahippocampus, left superior temporal pole, right insula, and right orbitofrontal cortex. The renderings were created using the AAL3 atlas and visualized in DSI Studio.

In addition, two behaviorally relevant SC-dFC group differences emerged. The effective connection from the left parahippocampus to the left superior temporal pole was stronger in gamers (*p* = 0.034) shown in Figure 3a(ii), and was negatively correlated with response times (*r* = –0.36, *p* = 0.022) shown in Figure 2b(xii). In contrast, the effective connection from the right insula to the right posterior orbital cortex was stronger in non-gamers (*p* = 0.046) shown in Figure 3a(iii), and was positively correlated with response times (*r* = 0.36, *p* = 0.025) demonstrated in Figure 2b(xvi). Figure 3b provides a visual representation of these connections in gamers and non-gamers, rendered using DSI Studio.

### 2.4. SC-FC Graph-Theoretic Network Analysis

After applying structural connectivity (SC) constraints to the functional connectivity (FC) data, we retained the top 95% of the strongest connections to construct binarized SC-FC graphs for network analysis. This threshold maximized the characterization of the SC-FC network while maintaining sparsity. At the global level, network measures, including characteristic path length, assortativity, and global efficiency, did not significantly differ between gamers and non-gamers.

To further investigate topological differences, we examined local graph-theoretic metrics, specifically local efficiency and node degree. Local efficiency reflects regional integration by measuring how effectively information is exchanged among a node’s immediate neighbors if the node itself is removed. Node degree, a local measure of centrality based on how many direct links a node has to other regions in the network, reflects the extent to which a region participates in the SC-FC network.

#### 2.4.1 SC-FC Local Efficiency & Node Degree Significant Differences

Gamers exhibited significantly greater local efficiency in the right middle occipital gyrus (*p* = 0.02) and right supramarginal gyrus (*p* = 0.047), suggesting stronger localized integration within dorsal visual and parietal circuits. In contrast, non-gamers showed greater local efficiency in the left pallidum (*p* = 0.047), a subcortical region involved in motor regulation and reinforcement learning.

For node degree, gamers demonstrated significantly higher values in the right inferior frontal gyrus (triangular part) (*p* = 0.015), right insula (*p* = 0.017), and two subdivisions of the left anterior cingulate cortex subgenual (*p* = 0.028) and pregenual (*p* = 0.032). These are key nodes in the salience and cognitive control networks, supporting integration of internal state monitoring and goal-directed action. By contrast, non-gamers showed higher node degree in the left cerebellum 3 (*p* = 0.009) and left hippocampus (*p* = 0.047), reflecting greater centrality in circuits involved in motor coordination and memory-based retrieval. These results are summarized using violin plots, which show the group distributions in Figure 4a.

**Figure 4.**
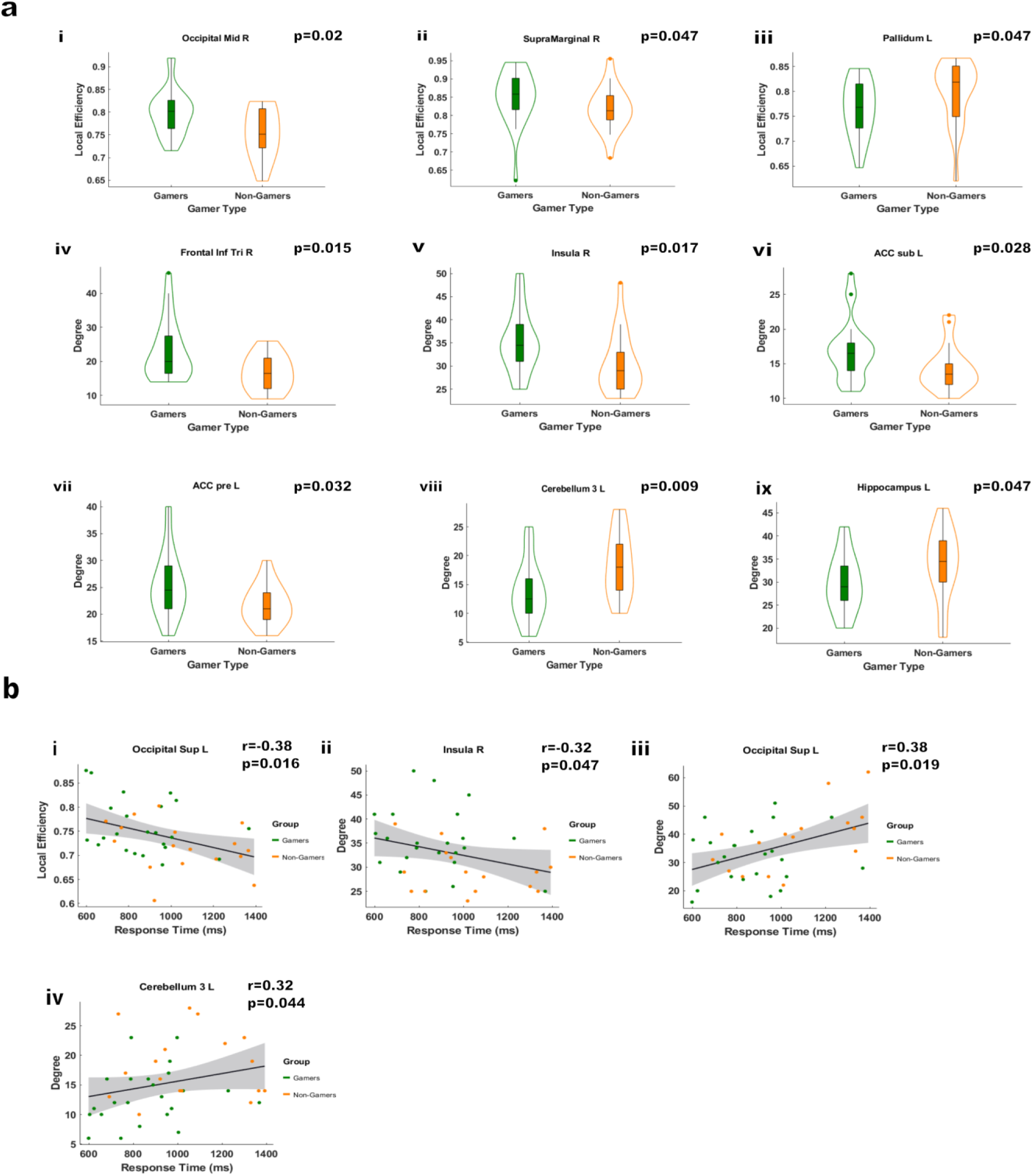
Group Differences in Binarized Functional Connectivity Network Metrics and Brain-Behavior. (a) Violin plots depicting group differences in binarized functional connectivity network metrics, including (i-iii) local efficiency and (iv-ix) node degree, for gamers and non-gamers. (b) Correlations between functional connectivity network metrics and response times. Negative correlations indicate an association with faster responses, while positive correlations reflect an association with slower responses.

#### 2.4.2 SC-FC Network Correlations with Response Times

Several SC-FC graph metrics were significantly correlated with response times across participants. Negative correlations were found for local efficiency in the left superior occipital gyrus (*r* = –0.38, *p* = 0.016) and node degree in the right insula (*r* = –0.32, *p* = 0.047), indicating that higher values were associated with faster responses. Positive correlations were observed for node degree in the left superior occipital gyrus (*r* = 0.38, *p* = 0.019) and left cerebellum 3 (*r* = 0.32, *p* = 0.044), where higher values were associated with slower responses. The brain–behavior correlations between SC-FC and RT are shown in Figure 4b.

### 2.5 SC-dFC Graph-Theoretic Network Analysis

For the SC-dFC analysis, we retained the top 10% of the strongest directed connections after applying SC constraints, accounting for the inherently sparser, asymmetric nature of dFC interactions. As in the SC-FC analysis, global network measures including characteristic path length, assortativity, density, and global efficiency did not significantly differ between groups. We examined local measures, specifically node degree and local efficiency, which are more sensitive to regional network properties and revealed significant differences between gamers and non-gamers.

#### 2.5.1 SC-dFC Local Efficiency & Node Degree Differences

Gamers exhibited significantly greater SC-dFC local efficiency in the right middle occipital gyrus (*p* = 0.026) and left precentral gyrus (*p* = 0.044), suggesting more integrated local processing within early visual and motor areas. Non-gamers, by contrast, had greater local efficiency in the left pallidum (*p* = 0.034) and vermis 4,5 (*p* = 0.047), indicating increased localized interaction within subcortical and cerebellar structures.

For node degree, gamers showed significantly higher values in multiple frontal and salience-related regions, including the left anterior cingulate (pregenual) (*p* = 0.008), right insula (*p* = 0.018), right rectus (*p* = 0.026), left superior medial frontal (*p* = 0.027), right inferior frontal (opercular) (*p* = 0.027), left posterior orbital (*p* = 0.045), and right inferior frontal (triangular) (*p* = 0.047). Non-gamers had significantly greater node degree in the left cerebellum 3 (*p* = 0.013) and left hippocampus (*p* = 0.031), consistent with stronger centrality in motor and memory-related regions. These results are summarized using violin plots, which show the group distributions in Figure 5a.

**Figure 5.**
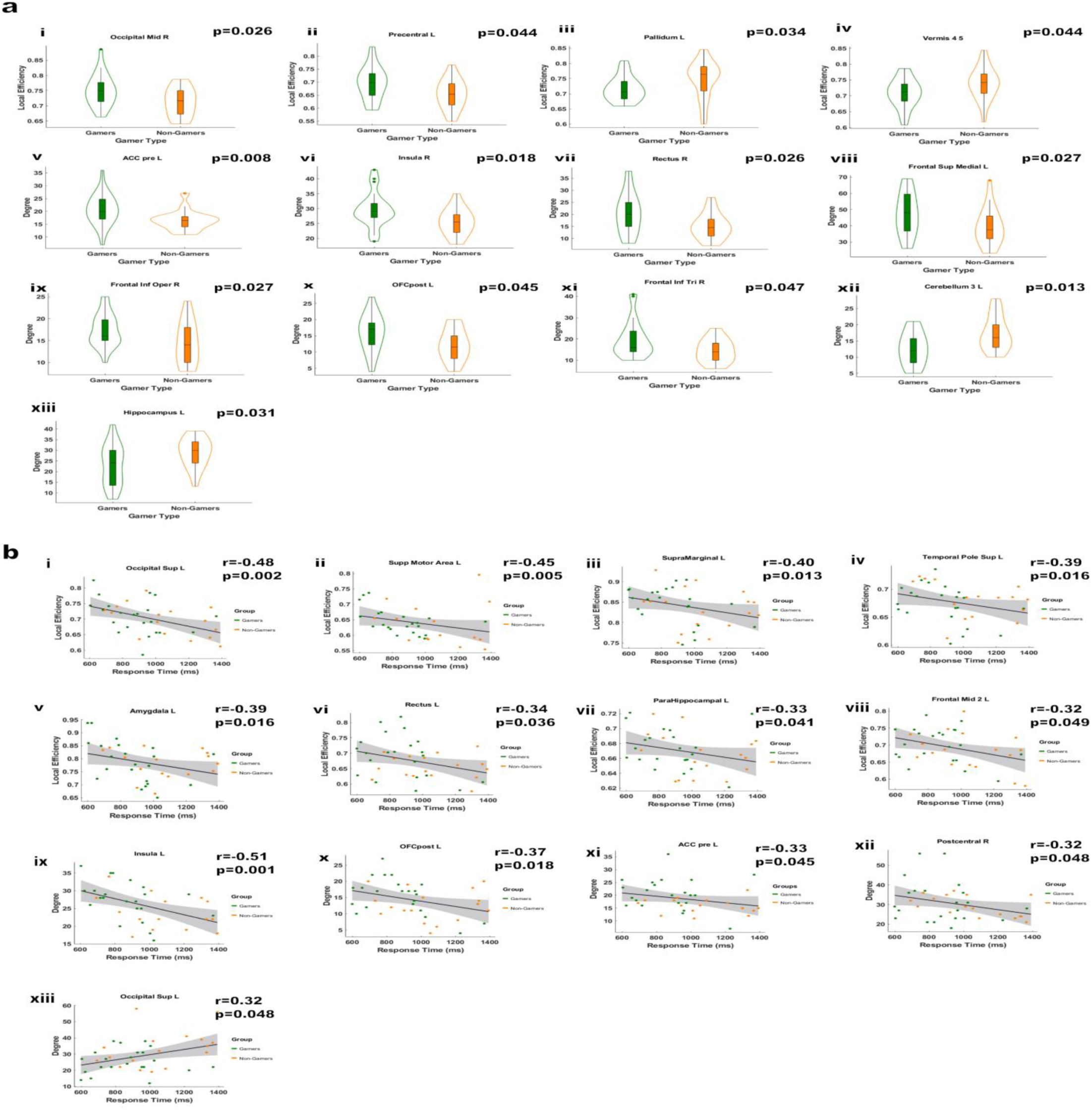
Group Differences in Binarized Directed Connectivity Network Metrics and Brain-Behavior. (a) Violin plots depicting group differences in binarized directed functional connectivity network metrics, including (i-vii) local efficiency and (ix-xii) node degree, for gamers and non-gamers.(b) Correlations between directed functional connectivity network metrics and response times. Negative correlations indicate an association with faster responses, while positive correlations reflect an association with slower responses.

#### 2.5.2 SC-dFC Network Correlations with Response Times

Both local efficiency and node degree SC-dFC were significantly correlated with response times (RT) across participants. We observed several negative correlations, indicating that higher graph values were associated with faster responses. For local efficiency, significant negative correlations were found in the left superior occipital (*r* = –0.48, *p* = 0.002), left supplementary motor area (*r* = –0.45, *p* = 0.005), left supramarginal (*r* = –0.40, *p* = 0.013), left superior temporal pole (*r* = –0.39, *p* = 0.016), left amygdala (*r* = –0.39, *p* = 0.016), left rectus (*r* = – 0.34, *p* = 0.036), left parahippocampus (*r* = –0.43, *p* = 0.006), and left middle frontal gyrus (*r* = –0.41, *p* = 0.009). For node degree, faster response times were associated with higher values in the left insula (*r* = –0.51, *p* = 0.001), left posterior orbital (*r* = –0.37, *p* = 0.018), left anterior cingulate (pregenual) (*r* = –0.33, *p* = 0.045), and right postcentral gyrus (*r* = –0.32, *p* = 0.048). Only one positive correlation was observed, with higher node degree in the left superior occipital gyrus associated with slower response times (*r* = 0.32, *p* = 0.048). The brain–behavior correlations between SC-dFC and RT are shown in Figure 5b.

## 3. Discussion

The results of this analysis provide compelling evidence that long-term action video game playing induces neuroplastic changes in structurally constrained functional and directed connectivity, leading to more efficient decision-making. These findings reveal distinct differences between gamers and non-gamers in connectivity patterns, brain-behavior relationships, and local network properties, suggesting a fundamental shift in visuomotor processing and decision-making strategies.

### 3.1 Structurally Constrained Functional Connectivity Profiles

#### 3.1.1 SC-FC Gamer and Non-Gamer Connectivity Patterns

Gamers exhibited a significantly greater number of enhanced (*p* < 0.05) SC-FC connections than non-gamers (278 ± 17 vs. 220 ± 15, *Z* = 2.60, *p* < 0.01). The SC-informed FC connectivity matrix shown in Figure 1a revealed more prominent connectivity shifts in favor of gamers across occipital-limbic, occipital-parietal, frontal-limbic, and frontal-parietal pathways. This pattern is consistent with the predictions of Cognitive Resource Reallocation (CRR), suggesting that video game experience enhances neural synchrony within circuits that support visual processing, attentional control, visuomotor integration, flexible action selection, and efficient decision-making under time pressure.

In the occipital-limbic pathway, increased SC-FC suggests stronger synchronization between regions involved in salience detection and early visual processing, potentially facilitating more effective extraction of task-relevant visual cues^46^. Enhanced occipital-parietal connectivity supports the integration of spatial and motion cues necessary for tracking object trajectories, indicating a greater reliance on endogenous attentional mechanisms to guide action ^47, 48^. Strengthened frontal-limbic connections reflect improved integration of executive and affective signals relevant to attentional control and adaptive behavior ^49, 50^. Finally, increased frontal-parietal coupling aligns with enhanced intentional action planning and selection during decision-making tasks, consistent with prior models of visuomotor coordination ^51^.

In contrast, non-gamers exhibited stronger SC-FC connectivity between frontal and occipital regions, suggesting a greater reliance on executive-visual synchrony rather than the anticipatory visuomotor response selection observed in gamers ^52^. Additionally, greater intra-cerebellar connectivity in non-gamers indicates a heavier reliance on feedback-driven motor adjustments, given the cerebellum’s known role in motor modulation^53^, which likely reflects a compensatory mechanism for less adaptive top-down motor planning and response execution. These findings suggest that non-gamers’ visuomotor processing strategies are less optimized, characterized by broader, more reactive, back-and-forth engagement between the visual, executive, and motor correction systems rather than the targeted, feedforward, adaptive response selection characterized by SC-FC connectivity patterns found in gamers.

#### 3.1.2 SC-dFC Gamer and Non-Gamer Connectivity Patterns

Non-gamers exhibited a greater total number of significantly stronger (*p* < 0.05) SC-dFC connections (313 ± 18 vs. 249 ± 16, *Z* = 2.70, *p* < 0.01) shown by the SC-dFC connectivity matrix in Figure 1b, suggesting a greater need for directed interactions to support their visuomotor decision-making. In contrast, gamers displayed more frontal-occipital and frontal-parietal SC-dFC connections, indicating a shift toward more targeted signaling between executive, visual, and motor regions, whereas non-gamers seem to rely more on broader, frontal-occipital engagement. Additionally, non-gamers exhibited significantly greater intra-cerebellar SC-dFC interactions, reinforcing their reliance on corrective motor adjustments rather than anticipatory control mechanisms.

### 3.2 SC-FC and SC-dFC Brain–Behavior Correlations

The following interpretations are grounded in well-established canonical neural anatomy and physiology of the involved brain regions, where functional roles are less well established in the literature.

#### 3.2.1 SC-FC Correlations with Response Times

Stronger connectivity between occipital, cerebellar, and multimodal sensory regions was associated with faster response times (RTs), suggesting that these pathways facilitate efficient visuomotor processing. This pattern is demonstrated clearly in Figure 2a. For example, connectivity between the left inferior temporal gyrus and the left cerebellum Crus I (*r* = –0.40, *p* = 0.012) suggests that motor planning and control processes, synchronized with object recognition (such as identifying moving target dots), facilitate faster decision-making and response execution. Similarly, connectivity between the right cerebellum Crus I and the right lingual gyrus (*r* = –0.38, *p* = 0.016) implies that visual scene processing paired with anticipatory motor planning plays a key role in rapid response execution.

Connectivity between the left insula and the left superior temporal gyrus (*r* = –0.37, *p* = 0.019) modulates interoceptive attention to auditory stimuli, effectively gatekeeping salient auditory information from executive engagement and optimizing cognitive resources for efficient visuomotor decision-making. Additional connectivity between the right calcarine and the left cuneus *(r* = –0.33, *p* = 0.039), as well as between the left cuneus and left middle occipital gyrus (*r* = –0.35, *p* = 0.029), indicates that enhanced early-stage visual processing supports rapid extraction of motion cues and enables quicker decision-making.

Conversely, stronger SC-FC connectivity in memory-related and feedback-driven motor regions has a positive correlation with RT, which is tracked with slower responses. This pattern suggests a reliance on deliberative processing rather than real-time visuomotor integration. For instance, connectivity between the left hippocampus and the left parahippocampus (*r* = 0.36, *p* = 0.021) points to an antagonism between scene-specific spatial configuration and object-in-place cognitive mapping, which may slow decision-making.

Similarly, stronger connectivity between the left cerebellum Crus I and the left vermis 4,5 (*r* = 0.38, *p* = 0.015) indicates increased reliance on corrective motor feedback, which could prolong response execution.

#### 3.2.2 SC-dFC Correlations with Response Times

To investigate how structurally constrained directed functional connectivity (SC-dFC) influences response time (RT), we assessed correlations between connectivity strength and RT across all participants, as depicted in Figure 2b.

Several SC-dFC connections were significantly associated with faster response times. For instance, directed connectivity from the right anterior cingulate (subgenual) to the right anterior cingulate (supracallosal) (*r* = – 0.51, *p* = 0.0009) was linked to urgency-driven response selection. This pathway is a major constituent of the dorsal attention network and likely serves as a high-priority signal that prompts executive systems to initiate rapid decision-making.

SC-dFC from the left middle temporal gyrus to the left lingual gyrus (*r* = –0.46, *p* = 0.003) supports the integration of high-level visual processing, such as object recognition, given the middle temporal gyrus’s proximity to the ventral stream, with color discrimination in the lingual gyrus. This integration may support rapid discrimination of color-based target dots from distractors during the sensory accumulation stage of a visuomotor decision.

Interactions from the left rolandic operculum to the left insula (*r* = –0.46, *p* = 0.003) and from the left insula to the left superior temporal gyrus (*r* = –0.46, *p* = 0.003) taken together suggest enhanced modulation of interoceptive attention to salient stimuli^46^. These pathways likely act to gate salient auditory information away from executive resources, facilitating scanner noise to be more of a persistent background feature than a salient distraction. Additionally, SC-dFC from the left superior temporal gyrus to the left rolandic operculum (*r* = – 0.44, *p* = 0.004) further supports streamlined sensory integration, likely under insular modulation, facilitating faster RT. Furthermore, based on known physiology, this loop may reflect interoceptive signaling prompting the retrieval or prioritization of high-level sensory information to guide an imminent motor response ^54^.

Connectivity from the left putamen to the left insula (*r* = –0.45, *p* = 0.005) and from the left putamen to the left superior frontal gyrus (*r* = –0.41, *p* = 0.009) highlights the putamen’s role in motor preparation and control^55^, suggesting that basal ganglia–insular circuits support the rapid coordination of motor action under time pressure.

Fronto-insular interactions were also predictive of faster response times. Directed signaling from the right inferior frontal gyrus (triangularis) to the right insula (*r* = –0.37, *p* = 0.019) suggests unconscious perceptual priming and enhanced attentional control ^56^. Likewise, connectivity from the right inferior frontal gyrus (orbital) to the right rolandic operculum (*r* = –0.37, *p* = 0.021) likely reflects a goal-directed control mechanism that bridges the brain’s interoceptive goal-directed map, such as the intention to make the correct decision, with voluntary motor execution of finger movement. ^54, 57^

Inter-parietal connections also tracked with faster responses. Directed signaling from the left supramarginal gyrus to the left superior parietal lobule (*r* = –0.37, *p* = 0.019), and from the left inferior parietal lobule to the left superior parietal lobule (*r* = –0.36, *p* = 0.022), supports visuospatial attention, motor planning, and sensorimotor integration. These dorsal stream pathways likely enhance rapid action selection by engaging the dorsal attention network.

SC-dFC from the right medial orbital frontal gyrus to the right rectus gyrus (*r* = –0.37, *p* = 0.022) may reflect goal-directed control over response selection. Although the precise cognitive role of the gyrus rectus remains under investigation, prior work suggests its involvement in value-based decision-making and executive control^58, 59^.

A SC-dFC connection from the left parahippocampal gyrus to the left superior temporal pole (*r* = –0.36, *p* = 0.022) supports the integration of scene-specific contextual information, which may facilitate quicker decisions by rapidly resolving target–distractor dynamics in complex visual environments by more readily integrating relative motion of the target compared to the distractor. Finally, signaling from the right insula to the right inferior frontal operculum (*r* = –0.35, *p* = 0.026) suggests close coordination between interoceptive and motor regions^60^, further reinforcing the importance of insular modulation in facilitating fast, goal-oriented actions.

One SC-dFC connection had a positive correlation and was significantly associated with slower response times, namely, the SC-dFC interaction from the right insula to the right posterior orbitofrontal cortex (*r* = 0.36, *p* = 0.025). Given the insula’s role in interoception and the orbitofrontal cortex’s function in decision inhibition and uncertainty evaluation, this connection may reflect a shift toward internal state monitoring and deliberative control, which slows down response execution ^58, 61, 62^.

### 3.3 Behavioral Correlates of SC-FC and SC-dFC Group Differences

We observed group-level differences in SC-FC and SC-dFC patterns between gamers and non-gamers that tracked with RT. Non-gamers exhibited stronger connectivity between the left middle temporal gyrus and left inferior temporal gyrus (*p* = 0.002) and showed a positive correlation with response times (*r* = 0.37, *p* = 0.018), suggesting a greater reliance on detailed object recognition before committing to a decision^63–65^. In contrast, gamers exhibited stronger connectivity from the left parahippocampal gyrus to the left superior temporal pole (*p* = 0.034) and was negatively correlated with response times (*r* = –0.36, *p* = 0.022). The parahippocampus is crucial for spatial scene processing ^66^, while the superior temporal pole is known as a convergent hub of high-level sensory information and perhaps of high-level information convergence in general ^67^.

Stronger SC-dFC from the parahippocampus to the superior temporal pole in gamers suggests greater integration of scene-specific contextual information, likely allowing for more efficient decision-making by more readily facilitating the rapid integration of the relative motion of target dots compared to distractors. Rather than solely relying on detailed object recognition, gamers place greater emphasis on the broader spatial and contextual relevance of the scene to guide their actions compared to non-gamers. This shift from object-based analysis to greater context-driven reasoning would contribute to greater decision efficiency in dynamic environments, reinforcing that long-term action video game playing experience leads to enhancements in adaptive visuomotor processing.

Additionally, non-gamers demonstrated stronger SC-dFC from the right insula to the right posterior orbitofrontal cortex (OFC) (*p* = 0.046) and had a positive correlation with RT (*r* = 0.36, *p* = 0.025). The right insula plays a key role in internal state monitoring and uncertainty assessment^61^. The posterior OFC contributes to evaluating outcomes and creating cognitive maps to navigate goal-directed behavior, such as the goal of picking the correct direction that target dots are moving in a visuomotor decision ^57^. Stronger connectivity involving these regions in non-gamers likely reflects a heightened emphasis on progressively reducing uncertainty before committing to a goal-directed decision of selecting the correct direction the target dots were moving, resulting in longer stimulus evaluation times at the expense of RT. This aligns with the well-established speed-accuracy tradeoffs in visuomotor decision-making, where prioritizing certainty and deliberation comes at the cost of slower response times ^68, 69^. In contrast, gamers likely engage in more real-time error correction to maintain accuracy and more effectively address uncertainty earlier in the decision-making process, enabling less reliance on feedback-driven inter-cerebellar corrections. This is supported by a shift in visuomotor decision strategy that more readily incorporates integration of scene-relevant information provided by their enhanced parahippocampal → superior temporal pole connectivity.

Together, these findings support the CRR hypothesis, demonstrating that long-term AVG experience plausibly promotes the reallocation of neural resources toward context-sensitive circuits that enable rapid, adaptive visuomotor decision-making. In contrast, non-gamers appear to rely more heavily on evaluative and uncertainty-monitoring systems that prioritize accuracy at the cost of speed. This divergence reflects distinct visuomotor decision-making strategies between gamers and non-gamers.

### 3.4 SC-FC Local Efficiency & Node Degree

For the undirected graph-theoretic network analysis, we applied a 95% threshold to binarize the SC-FC data, retaining the top 95% of the strongest connections after applying tractography constraints. This approach was chosen to capture as much of the structural network as possible while ensuring that only valid, non-spurious connections were included. As before, the following interpretations are grounded in canonical neural anatomy and physiology of the involved brain regions, with references provided where functional roles are less well established in the literature.

#### 3.4.1 Local Efficiency Group Differences

Graph-theoretic analysis revealed significant group-level differences in local efficiency, highlighting distinct patterns of network integration between gamers and non-gamers displayed in Figure 4a. Gamers exhibited significantly greater local efficiency in the right middle occipital gyrus (MOG) (*p* = 0.02). The right MOG plays a central role in integrating visual input with egocentric spatial orientation and processing spatial information, supporting visuomotor coordination ^70^. Increased local efficiency in this region likely facilitates low-latency visual processing and rapid motion tracking, advantages that are particularly beneficial in fast-paced visuomotor decision-making tasks requiring dynamic scene integration.

Gamers also demonstrated greater local efficiency in the right supramarginal gyrus (*p* = 0.047), a key node within the ventral attention network. This may reflect enhanced reorienting capacity, enabling more efficient shifts of attention to salient visual cues in dynamic environments such as action video games.

In contrast, non-gamers exhibited significantly greater local efficiency in the left pallidum (*p* = 0.047), a basal ganglia structure implicated in regulating voluntary movement and motor inhibition. This finding suggests a greater reliance on response inhibition mechanisms among non-gamers, which may contribute to slower, more deliberative decision-making strategies characterized by increased uncertainty monitoring.

#### 3.4.2 Node Degree Differences

Significant group differences were also observed in node degree, further elucidating the network-level reorganization associated with long-term gaming experience. Gamers exhibited higher node degree in several functionally relevant regions. These included the right inferior frontal gyrus triangularis (*p* = 0.015), a region central to executive control, unconscious perceptual priming, and information processing ^56^. Increased node degree here may reflect heightened readiness for rapid stimulus-response mapping and rule-based action selection.

Gamers also showed a greater degree in the right insula (*p* = 0.017), a salience network hub responsible for integrating sensory inputs and modulating attentional and decision-making processes. Elevated node degree was also observed in two subregions of the anterior cingulate cortex (ACC). In the subgenual ACC (*p* = 0.028), this increase is linked to urgency and affectively driven decisions, while in the pregenual ACC (*p* = 0.032), it reflects involvement in conflict monitoring and the adjustment of cognitive strategies in response to prediction errors. Together, these findings suggest that gamers leverage a more dynamically responsive network configuration that emphasizes anticipatory control and efficient adaptation to environmental demands.

In contrast, non-gamers exhibited higher node degree in regions associated with object-in-place cognitive mapping and feedback-driven motor regulation. Specifically, the left hippocampus (*p* = 0.047), a key structure for spatial memory and contextual mapping, showed greater centrality, indicating a strategy that relies more heavily on object-based spatial reasoning. Additionally, increased node degree was observed in the left cerebellar lobule 3 (*p* = 0.009), which is involved in motor feedback correction. This suggests that non-gamers depend more on feedback-driven motor adjustments, potentially resulting in slower response execution due to ongoing corrective processes rather than optimized feedforward planning.

#### 3.4.3 SC-FC Local Efficiency and Node Degree Correlations with Response Time

Stronger SC-FC local efficiency and node degree in visual and attentional regions were significantly associated with faster response times (RT), supporting their role in efficient visuomotor processing as shown in Figure 4b. Higher SC-FC local efficiency in the left superior occipital cortex (*r* = –0.38, *p* = 0.016) was associated with enhanced early visual processing, likely facilitating more efficient motion cue extraction and rapid response execution. Higher SC-FC node degree in the right insula (*r* = –0.32, *p* = 0.047), which was also significantly elevated in gamers (*p* = 0.017), was associated with faster response times. This aligns with the insula’s role in salience detection and adaptive attentional control, supporting the enhanced decision speed observed in gamers.

Conversely, stronger SC-FC node degree in early visual and feedback-driven motor regions was associated with slower response times, suggesting that an over-reliance on early-stage perceptual processing or corrective motor feedback may introduce inefficiencies. Specifically, higher node degree in the left superior occipital cortex (*r* = 0.38, *p* = 0.019) may reflect diffused or redundant spatial information processing that burdens downstream decision mechanisms. Similarly, increased node degree in the left cerebellar lobule 3 (*r* = 0.32, *p* = 0.044), which was significantly higher in non-gamers (*p* = 0.009), likely indicates greater reliance on feedback-based motor corrections, potentially delaying response execution due to slower, corrective adjustments during deliberation.

Taken together, these findings align with the CRR hypothesis by demonstrating that more efficient SC-FC integration in gamers supports a shift toward feedforward-dominant processing. This would enable more rapid visuomotor decisions based on salient visual cues. In contrast, non-gamers appear to rely more heavily on slower, feedback-dependent strategies, which impose greater cognitive load and contribute to delayed motor responses.

### 3.5 SC-dFC Local Efficiency and Node Degree Differences

SC-dFC is inherently asymmetric, meaning that a connection from source A to target B does not imply the same or a similar connection from B to A. This asymmetry results in dFC networks exhibiting greater variability and a sparser structure compared to FC networks.

To address this sparsity, we applied a 10% threshold to binarize the SC-dFC network, retaining only the top 10% of the strongest and most reliable directed interactions. This threshold minimized the influence of invalid or spurious connections, while preserving as many effective connections as possible. By doing so, we ensured that the global density of the SC-dFC network remained comparable to that of the SC-FC network within the same participant. This thresholding strategy aligned with our goal of maintaining as many meaningful connections as possible for an accurate characterization of the SC-dFC network while excluding invalid and spurious ones. The following interpretations are grounded in canonical neural anatomy and physiology of the involved brain regions, with references provided where functional roles are less well established in the literature.

#### 3.5.1 SC-dFC Local Efficiency Differences

Gamers demonstrated significantly greater SC-dFC local efficiency in two regions. First, the right middle occipital gyrus (*p* = 0.026), a region involved in motion perception and spatial processing, exhibited enhanced efficiency. This region likely serves as a transitional node linking early-stage visual areas, such as the superior occipital and calcarine cortices, to higher-order visuomotor networks. Greater local efficiency in this region suggests more effective directional information transfer, potentially enabling faster, low-latency visual processing of spatial information that is advantageous for rapid response execution ^70^. Second, gamers also showed increased local efficiency in the left precentral gyrus (*p* = 0.044), a primary motor area. This enhancement indicates improved sensorimotor coupling, allowing for more direct and efficient communication between visual input and motor output pathways.

In contrast, non-gamers exhibited significantly greater SC-dFC local efficiency in two different regions. The left pallidum (*p* = 0.034), a basal ganglia structure critical for proprioception and habitual motor actions, showed higher efficiency, suggesting a greater reliance on pre-learned or routine motor responses rather than executive planning. Additionally, the vermis 4,5 (*p* = 0.047), part of the cerebellum involved in error correction and feedback-driven motor coordination, also exhibited greater efficiency in non-gamers. This pattern implies a heavier dependence on corrective strategies, which may introduce temporal delays in response execution during time-sensitive tasks.

#### 3.5.2 SC-dFC Node Degree Differences

Examining SC-dFC node degree revealed that gamers exhibited significantly higher connectivity in several regions associated with executive control and adaptive decision-making. The left anterior cingulate cortex pregenual (*p* = 0.008), known for its role in performance monitoring and adaptive control, showed enhanced node degree, suggesting greater integration of cognitive control mechanisms to support rapid response selection. The right insula (*p* = 0.018), a core node of the salience network, also displayed increased connectivity, reinforcing the idea that gamers are more adept at optimizing sensory-cognitive interactions for stimulus-driven decision-making.

Additional regions showing increased node degree in gamers included the right rectus gyrus (*p* = 0.026), thought to contribute to motivational behavior and reward-based decision-making ^58, 59^ and the left superior medial frontal gyrus (*p* = 0.027), which is associated with higher-order motor planning and strategic control. Furthermore, the right inferior frontal gyrus, both opercular (*p* = 0.027) and triangular (*p* = 0.047) parts, showed elevated node degree, pointing to more effective executive suppression of irrelevant signals and potentially enhanced unconscious priming. Finally, the left posterior orbital cortex (*p* = 0.045), involved in response evaluation and strategic adjustment, also demonstrated increased connectivity in gamers.

In contrast, non-gamers showed significantly higher node degree in regions associated with memory-guided and feedback-driven motor processing. These included the left hippocampus (*p* = 0.031), implicated in spatial memory and object-in-place mapping, and the left cerebellum lobule 3 (*p* = 0.013), which plays a role in corrective motor control. These patterns suggest that non-gamers rely more on feedback-dependent and memory-guided decision-making strategies rather than streamlined, stimulus-response circuits.

Collectively, these results support the notion that gamers exhibit more feedforward-driven visuomotor integration and executive control, enabling rapid response selection without the need for extensive uncertainty reduction. This is evidenced by greater SC-dFC local efficiency in key visual and motor integration areas, as well as increased node degree in salience and control-related regions. In contrast, non-gamers appear to rely more on habitual, feedback-corrective strategies involving memory and cerebellar coordination, which may introduce delays in rapid decision-making contexts.

#### 3.5.3 SC-dFC Efficiency and Node Degree Correlations with Response Times

Spearman correlations revealed that enhanced SC-dFC local efficiency in several regions was significantly associated with faster response times. Notably, efficiency in the left superior occipital cortex (*r* = –0.48, *p* = 0.002) supported enhanced early-stage visual processing and motion cue extraction, while the left supplementary motor area (*r* = –0.45, *p* = 0.005) contributed to motor planning and rapid visuomotor responses. The left parahippocampal gyrus (*r* = –0.43, *p* = 0.006), known for scene-based spatial memory, and the left supramarginal gyrus (*r* = –0.40, *p* = 0.013), involved in sensorimotor integration and attentional shifts, were also significantly correlated with response speed. Additionally, the left superior temporal pole (*r* = –0.39, *p* = 0.016), thought to serve as a hub for perceptual integration ^67^, and the left amygdala (*r* = –0.39, *p* = 0.016), involved in emotional salience processing, both showed significant negative correlations with response time.

Finally, local efficiency in the left rectus (*r* = –0.34, *p* = 0.036) was associated with faster response times, consistent with its role in motivational orientation and goal-directed behavior^58, 59^.

In terms of node degree, several regions demonstrated significant associations with faster response times. These included the left insula (*r* = –0.51, *p* = 0.001), a salience network hub that integrates sensory and interoceptive inputs and also showed significantly greater node degree in gamers (*p* = 0.017); the left posterior orbital cortex (*r* = –0.37, *p* = 0.018), involved in adaptive decision-making and similarly elevated in gamers (*p* = 0.045); and the left anterior cingulate cortex pregenual (*r* = –0.33, *p* = 0.045), which supports performance monitoring and adaptive control and was also higher in gamers (*p* = 0.008). The right postcentral gyrus (*r* = –0.32, *p* = 0.048), associated with sensorimotor integration and proprioception, also showed a significant behavioral correlation, although this region did not show a group difference in node degree.

Interestingly, a higher node degree in the left superior occipital gyrus was associated with slower response times (*r* = 0.32, *p* = 0.048), suggesting a potential tradeoff in information distribution. From the perspective of CRR, this may reflect a diffusion of vital visual information of object trajectory across multiple processing routes, analogous to current splitting in a parallel electrical circuit, which may slow decision-making.

Altogether, these findings underscore the role of enhanced SC-dFC efficiency and node degree in visuomotor and executive regions as contributors to faster response execution in gamers. Enhanced efficiency in early visual and sensorimotor regions, along with greater connectivity in salience and control hubs, appears to facilitate rapid stimulus-response transformations. These results align with the CRR hypothesis, suggesting that gamers optimize directional communication along more effective pathways, which would minimize reliance on cerebellar feedback loops and reduce response latency. In contrast, non-gamers’ greater reliance on memory and feedback-driven control systems may underlie slower, more deliberative response strategies.

### 3.6 Integration with Prior Research

This study builds on prior research into video game-induced neuroplasticity by further contextualizing previously reported findings from this dataset and demonstrating how structurally constrained functional and directed connectivity changes contribute to enhanced visuomotor decision-making response time (RT) ^28, 32^. Several key regions previously identified as critical for video game-related neural enhancements reappear in the current analysis, particularly in the domains of visuomotor processing and attentional coordination.

One notable example is the right lingual gyrus, which in earlier work showed significant group differences in BOLD activation between gamers and non-gamers. In the present analysis, this region exhibited stronger SC-FC connectivity with the cerebellum (*r* = –0.38, *p* = 0.016), a relationship that also correlated with faster RT. This finding reinforces the lingual gyrus’s role in supporting rapid visual-motor transformations ^32^.

Additionally, improvements in connectivity between the dorsal attention network (DAN) and the salience network (SN) were observed in gamers, surviving Bonferroni correction (*p* < 0.05). These enhancements suggest more efficient attentional coordination and flexible network switching, likely enabling gamers to focus more effectively on task-relevant stimuli^32^. Expanding on these results, the current findings indicate that gamers differ from non-gamers not only in top-down DAN→SN attention allocation but also in how they process motion information. Specifically, the data support the idea that, through long-term engagement with action video games, cognitive resources are gradually reallocated toward more optimized neural pathways that incorporate scene-specific visual information, particularly relative motion cues, which in turn improve RT during visuomotor decision-making tasks.

A central region in this dynamic is the supracallosal (dorsal) anterior cingulate cortex (ACC), a key node within the salience network involved in top-down attention and internal conflict monitoring between competing motor plans. In the current study, gamers exhibited significantly higher SC-dFC node degree in both the subgenual (*p* = 0.028) and pregenual (*p* = 0.008) ACC, suggesting greater centrality in circuits involved in urgency signaling, adaptive control, and performance monitoring. Notably, node degree in the left pregenual ACC was also correlated with faster RT (*r* = –0.33, *p* = 0.045), consistent with its role in resolving action conflicts under pressure and optimizing behavioral responses.

Although the subgenual ACC did not correlate with RT, its elevated connectivity likely reflects affective modulation, heightening motivational salience and time-sensitive urgency signaling to the suprarcallosal ACC. These two regions appear to work in tandem, with the subgenual ACC driving arousal and the supracallosal ACC coordinating rapid executive responses. Supporting this view, stronger directed connectivity from the right subgenual to right supracallosal ACC was significantly associated with faster RT (r = –0.51, p = 0.0009), suggesting that a streamlined affective-to-executive signaling pathway is a strong indicator of decision-making RT.

Additional SC-dFC connections involving pregenual and supracallosal ACC regions also tracked with faster RT. These included connections from the right pregenual ACC to the left pregenual ACC (*r* = –0.33, *p* = 0.036), from the left pregenual ACC to the right supracallosal ACC (*r* = –0.38, *p* = 0.0169), and from the right pregenual ACC to the right supracallosal ACC (*r* = –0.33, p = 0.038).

Our results also reinforce earlier findings, which showed that functional connectivity in the left dorsal stream is enhanced in gamers^38^ (Holm-Bonferroni corrected, p < 0.05), with a significant correlation to faster response times (*r* = –0.41; Holm-Bonferroni corrected, *p* < 0.05). Specifically, gamers exhibited increased FC in the left superior occipital gyrus (L SOG) and superior parietal lobule (L SPL), core dorsal stream regions essential for visuomotor integration. Faster response times were significantly associated with higher local efficiency in both SC-FC (*r* = –0.38, *p* = 0.016) and SC-dFC (*r* = –0.48, *p* = 0.002) within the left superior occipital gyrus (L SOG), supporting the interpretation that enhanced dorsal stream function in gamers facilitates more efficient trajectory estimation and visuomotor integration.

Interestingly, however, greater SC-FC node degree in the L SOG had a positive correlation with RT (*r* = 0.38, *p* = 0.019), suggesting that excessive reliance on early-stage visual processing may lead to a less efficient visuomotor strategy—analogous to splitting current across too many paths in a parallel circuit.

Collectively, these findings lend further support to the CRR hypothesis, which posits that long-term action video game experience drives a reallocation of cognitive resources toward more efficient visuomotor decision-making circuits. By reinforcing salience detection and motion tracking systems, AVG experience would enable rapid, accurate responses in high-pressure environments. These results underscore the potential role of adaptive neural processing and network refinements as key mechanisms that underlie AVG-induced neuroplasticity.

### 3.7 Evaluation of the CRR Hypothesis

The goal of this study was to formally evaluate the Cognitive Resource Reallocation (CRR) hypothesis as a plausible mechanistic explanation for the enhanced visuomotor decision-making observed in long-term action video game (AVG) players. CRR posits that sustained AVG engagement gradually reallocates cognitive resources toward more efficient, feedforward visuomotor pathways, favoring circuits that support rapid, goal-directed responses over slower, deliberative feedback loops. This reallocation is expected to manifest as anatomically plausible changes in both the structure and dynamics of neural networks, supporting more efficient visuomotor decision making through enhancements in visual processing, visuomotor integration, attentional control, and cognitive flexibility. To test this hypothesis, this study focuses on group-level differences that also showed significant correlations with response time (RT). These findings represent the most direct evidence with which to evaluate CRR as a candidate explanation of the behavioral advantage observed in gamers.

We observed group-level differences in SC-FC and SC-dFC that tracked with RT. For example, non-gamers exhibited stronger SC-FC between the left middle temporal gyrus and left inferior temporal gyrus (*p* = 0.002), a connection positively correlated with slower RTs (*r* = 0.36, *p* = 0.025). This suggests a decision-making strategy weighted toward detailed object recognition, potentially increasing stimulus evaluation time. By contrast, gamers showed stronger SC-dFC from the left parahippocampal gyrus to the left superior temporal pole (*p* = 0.034), a connection negatively correlated with RT (*r* = –0.36, *p* = 0.022). This pathway likely facilitates more efficient integration of scene-specific contextual information and relative motion, aligning with a context-driven decision-making strategy.

This interpretation is supported by greater SC-dFC local efficiency in both the left superior temporal pole (*p* = 0.029, *r* = –0.33) and the left parahippocampus (*p* = 0.033, *r* = –0.34), indicating more efficient local integration of scene-relevant information in gamers. While efficiency in these regions did not differ between groups, the fact that gamers demonstrated stronger SC-dFC between them, coupled with the negative RT correlation, supports a shift toward optimized visuospatial processing of salient scene-specific cues.

In contrast, non-gamers demonstrated stronger SC-dFC from the right insula to the right posterior orbitofrontal cortex (OFC) (*p* = 0.043), a connection positively correlated with RT (*r* = 0.36, *p* = 0.025), tracking with slower response times. The right insula is involved in interoception and uncertainty monitoring, while the posterior OFC supports outcome evaluation and decision inhibition. Stronger SC-dFC signaling from the right insula to the right posterior OFC likely reflects a heightened emphasis on internal deliberation and iterative uncertainty reduction in non-gamers, amplifying the speed-accuracy tradeoff by promoting accuracy at the cost of RT.

In line with this interpretation, gamers exhibited significantly higher SC-FC node degree in the right insula (*p* = 0.017), which was associated with faster RT, as indicated by its negative correlation (*r* = –0.32, *p* = 0.047).

While both groups appear to engage the right insula during visuomotor decision-making, gamers were shown to benefit more from its involvement due to greater network centrality, amplifying its role in rapid salience detection and adaptive interoceptive control. From this evidence, it is clear that gamers tend to leverage right insular engagement more effectively by tending to establish it as more central to the decision-making process to interoceptively monitor more regions, tending to devote less resources to support a directed connection to the posterior OFC, while non-gamers tend to recruit the right insula in such a way that aligns more closely with canonical speed-accuracy tradeoffs.

SC-FC and SC-dFC node degrees also revealed consistent enhancements in gamers. SC-FC node-degree of the left cerebellum lobule 3 tended to be greater in non-gamers (*p* = 0.009), and correlated with slower RTs (*r* = 0.32, *p* = 0.044), indicative of a more feedback-dependent motor correction strategy, once again aligning with a canonical speed-accuracy tradeoff. In contrast, the pregenual ACC had a higher SC-dFC node degree in gamers (*p* = 0.008) and was significantly correlated with faster RTs (*r* = –0.33, *p* = 0.045), suggesting a role in top-down attentional control and more effective resolution of internal conflict regarding competing motor plans.

Higher local efficiency in the left superior occipital gyrus (SOG) was associated with faster response times in both SC-FC (*r* = –0.38, *p* = 0.016) and SC-dFC (*r* = –0.48, *p* = 0.002). Although no group differences were observed in SOG efficiency, these results reinforce the dorsal stream’s critical role in visuomotor integration, particularly during the sensory accumulation phase of decision-making, where object motion and trajectory must be rapidly estimated and an imminent motor response is required. As shown in previous work, this system is functionally enhanced in gamers, underscoring the dorsal stream’s contribution to more efficient visuomotor decision-making^38^.

Taken together, the evidence from this study rejects the null hypothesis that CRR is not a plausible explanatory principle for the visuomotor decision-making advantage observed in gamers. Across SC-FC and SC-dFC modalities, local network properties, and response time correlations, we observed converging evidence that long-term AVG engagement reflects a reallocation of cognitive resources, which would induce neuroplastic refinements that support a more effective visuomotor decision-making strategy. Notably, no contradictions were observed within this dataset, providing unilateral support for CRR as a viable mechanistic account of experience-driven neuroplastic refinement associated with enhanced visuomotor decision-making in the context of AVG experience.

While CRR was found to be strongly supported as a plausible mechanistic explanation for enhanced visuomotor performance in gamers, it is important to consider what kinds of findings would have contradicted the hypothesis. CRR would be challenged if non-gamers showed stronger connectivity or network properties in task-relevant circuits that also predicted faster response times, especially if such markers were absent or weaker in gamers. It would also raise concerns if neural enhancements in gamers were limited to only part of the visuomotor decision process, while non-gamers showed stronger tuning in other equally relevant components. Another challenge to CRR would come from evidence that pre-existing individual differences, such as globally more efficient networks unrelated to task relevance, predispose individuals both to faster performance and a higher likelihood of gaming. This would suggest a selection effect rather than a plasticity-driven process.

Similarly, if non-gamers showed stronger brain–behavior correlations in task-relevant areas than gamers, that too would challenge CRR. However, none of these patterns were observed. Instead, we found consistent adaptations in gamers across task-relevant networks that closely track behavioral performance. These findings support cognitive resource reallocation as the mechanism underlying the observed neuroplastic changes.

### 3.8 Limitations

Our study recruited a sample of healthy young adults, allowing us to isolate the effects of long-term video game playing while minimizing confounds. However, this design choice also imposes certain limitations. Addressing these limitations in future studies will be beneficial to fully characterize the extent and generalizability of video game-induced cognitive and neural adaptations. First, the dataset used in this study captures a cross-sectional snapshot of individuals with long-term action video game experience. As such, it does not permit inferences about the rate of neuroplastic adaptation over time, limiting the ability to make direct causal claims.

Determining how these changes evolve would require longitudinal studies and clinical training interventions. Additionally, our gender distribution was not balanced between gamers and non-gamers, precluding direct analysis of gender-specific effects.

Although prior research suggests that action video game experience may mitigate cognitive decline in older adults ^71^, our study exclusively focused on young adults, leaving open the question of how the neuroplastic effects observed in this study extend across different age groups. While participants were recruited from university campuses with presumably similar educational backgrounds, we did not explicitly screen for education levels or cognitive ability, meaning we cannot establish direct correlations between baseline cognitive performance and task outcomes. Furthermore, although our sample size was sufficient for statistical analysis, future studies should expand sample diversity to enhance generalizability. Most importantly, greater statistical power would enable more sensitive testing of within-group brain–behavior correlations, making it possible to determine whether observed effects are driven primarily by gamers, non-gamers, or both. Therefore, larger cohorts would support a more granular analysis of individual group-specific differences and enable more robust comparisons across gaming subgenres^72^.

Furthermore, it would have been valuable to compare connectivity profiles for the moving dots task with rs-fMRI data^73^. This could have offered additional insight into how task-based network dynamics relate to intrinsic functional organization. However, resting-state scans were not collected as part of this study, limiting our ability to explore these relationships.

Methodologically, using white matter tractography to structurally constrain functional connectivity reduces false positives by limiting statistical comparisons to anatomically viable connections—i.e., those supported by known white matter tracts. Unlike unconstrained full-brain FC analyses, which test all possible pairwise combinations regardless of biological plausibility, considering SC-FC and SC-dFC improves neurobiological interpretability and dramatically reduces the total number of comparisons.

Specifically, both SC-FC and SC-dFC analyses were computed across 13,818 valid statistical comparisons (excluding NaNs and self-connections). Under a null model with α = 0.05, this would yield approximately 690 ± 26 false positives. In contrast, we observed substantially fewer: 498 for SC-FC (*Z* = –7.38, *p* = 1.53 × 10⁻¹³) and 592 for SC-dFC (*Z* = –3.77, *p* = 0.00016). The observed distributions deviate significantly from the null expectation, strongly indicating that these results reflect structured, behaviorally meaningful differences in connectivity rather than random noise.

The veracity of our results is further supported by consistent and interpretable violin plot distributions for node degree and local efficiency, which strengthens confidence in this framework. Our findings align with known neuroanatomical pathways and task-relevant brain systems, reinforcing the utility and validity of SC-based filtering for detecting meaningful group-level effects.

However, structural connectivity is inherently constrained by tractography’s parameter bounds—minimum fiber length, angular thresholds, and model resolution. This may result in the omission of short-range, sharply turning, or multi-synaptic pathways, especially within complex relay hubs like the thalamus. Moreover, SC-based filtering excludes connections between regions that are functionally coordinated but lack a directly detected structural edge.

Tractography remains an indirect approximation of true anatomical architecture. While we mitigate some of its limitations by using quantitative anisotropy (QA)–based tractography, which outperforms traditional FA in resolving crossing fibers and preserving directional specificity, no tractography method offers absolute anatomical ground truth. QA, based on q-space diffeomorphic reconstruction (QSDR), improves sensitivity to true axonal trajectories and bolsters anatomical plausibility. Nonetheless, tractography fidelity ultimately constrains the scope of SC-based connectivity.

While SC-FC and SC-dFC analyses offer anatomical grounding, they may miss distributed patterns of functional change that fall outside direct white matter pathways. Integrating this method with complementary data-driven approaches may offer a promising way to recover signals that may be overlooked by strict structural constraints.

Finally, functional ROI measures derived from fMRI are limited in temporal resolution and may fail to capture finer-grained dynamics such as fast oscillatory coupling or cross-network phase interactions. Since the task involved only a simple button-press response, our analyses focused on visuomotor decision-making, assuming that motor execution was not the primary source of group differences. However, future studies employing event-related fMRI, electroencephalography (EEG), or magnetoencephalography (MEG) could help distinguish decision-making processes from motor execution components.

### 3.9 Future Considerations

Our study provides a strong foundation for understanding how long-term action video game play reshapes brain connectivity, supporting more efficient visuomotor decision-making that enables faster response times without tradeoffs in accuracy. As the field evolves, future studies can build upon our findings by leveraging larger and more diverse datasets to enhance generalizability and refining methodological approaches to bolster future research standards in this field, ensuring continued rigor in experimental design, neuroimaging analysis, and behavioral assessment. Furthermore, a key improvement would be the inclusion of resting-state fMRI (rs-fMRI) data, which would have allowed us to examine baseline connectivity patterns without the task and determine whether gaming-related neuroplasticity extends beyond task-driven effects to broader network-level adaptations.

Additionally, parsing subgenre-specific effects to determine how different game mechanics influence neural adaptations, conducting more clinical and longitudinal video game training studies to track neuroplastic changes over time, and identifying critical windows for skill transfer. To maximize translational value, future work should prioritize clear brain-behavior relationships, using structured assessments across training intervals to capture the rate and progression of induced effects, and pairing the findings here with data-driven methods could help recover fine-grained connections, such as intra-thalamic connections, that were lost due to minimum length considerations in our tractography, providing a more comprehensive understanding of neural adaptations across all task-relevant brain regions.

Our work lays the groundwork for reframing video games not only as a tool for applied intervention and cognitive training but also as a research paradigm for inducing experience-driven neuroplasticity. By linking neural changes to observable improvements in cognitive performance, we can better understand how video games drive shifts in cognitive resources, enhancing attentional control, sensorimotor integration, and executive response selection. These insights extend far beyond gaming itself, offering valuable implications for cognitive training, rehabilitation, and skill acquisition in high-performance settings like surgery, aviation, and military operations.

The focus of future research should shift toward optimizing targeted video games, designed to deliberately influence specific cognitive functions. This can be validated using structurally constrained functional analysis, as we have done in our work, providing a robust framework for understanding the underlying neural mechanisms at play.

While video games are known to enhance aspects of cognitive performance, uncovering the neural mechanisms behind these changes offers even greater value. Identifying how the brain reallocates cognitive resources and reorganizes network connectivity to drive these effects is crucial. This isn’t just about validating psychological findings; it’s about explaining how the brain adapts and refines cognitive skills through long-term gameplay. A deeper understanding of these mechanisms could optimize training and rehabilitation interventions, tailoring gaming experiences to enhance specific cognitive functions such as attention, memory, or problem-solving.

Beyond enhancing visuomotor decision making in the case of AVGs, video games may also promote meaningful psychological and emotional adaptations. These effects represent a promising dimension of neuroadaptive gaming, with potential applications across both clinical and non-clinical populations. Emerging research suggests that gameplay may influence emotional regulation, social cognition, resilience, and identity development. Specific game mechanics and genres may differentially support these outcomes. Narratively-rich games can scaffold emotional learning and recovery through exposure to challenge, loss, and moral decision-making^74^. Games that simulate adversity and perseverance may offer therapeutic value for coping strategy development and psychological resilience^75^. Multiplayer and cooperative games can support the practice of social behaviors in low-stakes settings, potentially improving emotional expression, empathy, and interpersonal communication ^76^. Open-ended “sandbox” games that reward flexible thinking and exploration may facilitate cognitive flexibility and lateral thinking^77, 78^.

As the field of video game-based interventions continues to develop, it presents potential avenues for enhancing cognitive, emotional, and social well-being, with possible applications across both clinical and non-clinical populations. Understanding the underlying neural mechanisms driving these effects can help refine and improve interventions aimed at maximizing these outcomes by providing insights into how the brain allocates and reallocates cognitive resources to optimize performance on tasks requiring specific cognitive abilities to engage with and play these video games effectively. Ultimately, this knowledge may inform the design of tailored video games optimized for cognitive training and rehabilitation, supporting the reallocation of resources to neural networks involved in cognitive function, emotional regulation, and pro-social behavior.

### 3.10 Concluding Remarks

This study investigates how long-term action video game play could induce neuroplastic refinements that enhance visuomotor decision-making and its supporting cognitive functions. While prior research has established that gamers exhibit faster response times and cognitive benefits, the specific neural mechanisms underlying these improvements have remained unclear.

Our findings unilaterally support Cognitive Resource Reallocation (CRR) as the underlying principle governing gaming-related neuroplasticity, with no contradictions observed across any facet of our results. This shift reflects a transition from feedback-driven motor correction to anticipatory, feedforward processing, optimizing response efficiency and enhancing relative motion tracking between targets and distractors. These improvements are supported by strengthened connectivity in circuits involved in visual processing, visuomotor integration, attentional control, and cognitive flexibility, resulting in a more streamlined neural architecture for action selection in dynamic environments.

This conclusion, evidenced by SC-FC and SC-dFC connectivity analyses, graph-theoretic metrics, and behavioral correlations with response time, is consistent with CRR’s mechanistic account of how cognitive resources are selectively reallocated to task-relevant pathways. This reallocation leads to measurable behavioral advantages, including an average improvement of approximately 190 milliseconds in response time among gamers. By supporting CRR as a plausible explanatory framework for experience-dependent neuroplasticity, this work advances our understanding of how intense visuomotor engagement refines brain networks to support rapid, adaptive decision-making in fast-paced, graphically rich environments such as AVGs.

These findings provide a strong basis for further investigation, establishing video game play not only as a topic of interest to the cognitive training and rehabilitation community but also as a medium through which general organizing principles of neuroplastic adaptation may be more completely understood. Future research should further explore CRR as a general organizing framework in video game playing and in other forms of skill acquisition, cognitive training paradigms, and real-world visuomotor expertise, shedding light on the generalizability of experience-driven neuroplasticity across behavioral domains.

Video game research must move beyond descriptive studies and adopt more rigorous methodologies to uncover the causal mechanisms of neuroplastic adaptation that drive changes in cognitive function. It is essential to ensure that these adaptations are behaviorally relevant and demonstrate skill transfer through direct cognitive assessments and comparisons. Integrating structural and functional neuroimaging data could enhance our understanding of the neural mechanisms underlying these cognitive adaptations. If Cognitive Resource Reallocation (CRR) serves as a general organizing principle for experience-induced neuroplasticity, it could explain how mesoscale adaptations drive large-scale, brain-wide network reconfigurations, resulting in measurable cognitive and psychological benefits across both clinical and healthy populations. By integrating neuroscience, cognitive science, and interactive digital media, not only could video games serve as tools for skill development and rehabilitation, but also as an ecologically rich experimental paradigm for understanding and guiding experience-dependent neuroplasticity.

## 2. Materials and Methods

### 2.1. Participant Data

47 right-handed participants (28 gamers (4 female) and 19 non-gamers (12 female)) were recruited. Groups were age-matched (gamers = 20.6 ± 2.4 years; non-gamers = 19.9 ± 2.6 years). Participants who indicated playing 5 hours per week or more in one of four types of video game genres for the last two years were considered “gamers”. The four types of action video game genres considered were First-Person Shooter (FPS), Real-Time Strategy (RTS), Multiplayer Online Battle Arena (MOBA), and Battle Royale (BR). Participants who were classified as “non-gamers” in this study averaged less than 30 minutes per week in any video game over the last two years. A modified left–right moving-dot task was used to probe for differences in response time and accuracy between the cohorts^32^. The effective number of participants for the SC-FC and SC-TGC analysis was 42 total participants (24 gamers and 18 non-gamers), with 40 total participants (23 gamers and 17 non-gamers) for brain-behavior regression between SC-FC & SC-TGC measures with response time. To confirm eligibility and assign participants to the appropriate group, a questionnaire was administered assessing video game genre and play frequency over the past two years. All participants passed the Ishihara Test for Color Deficiency and completed informed consent and health screening forms before data collection. The study was approved by the Institutional Review Boards of Georgia State University and the Georgia Institute of Technology, both located in Atlanta, Georgia.

### 2.2. MRI Data

#### 2.2.1 Data Collection, Scanning & Tractography Protocols

Whole-brain structural and functional MR imaging was conducted on a 3T Siemens Magnetom Prisma MRI scanner (Siemens, Atlanta, GA, USA) at the joint Georgia State University and Georgia Institute of Technology Center for Advanced Brain Imaging, Atlanta, GA, USA. High-resolution anatomical images were acquired using a T1-MEMPRAGE scan sequence for voxel-based morphometry and anatomical reference. The acquisition parameters were as follows: TR = 2530 ms, TE1-4 = 1.69–7.27 ms, TI = 1260 ms, flip angle = 7°, and voxel size = 1 mm × 1 mm × 1 mm.

Diffusion-weighted imaging (DWI) data were collected using a multi-shell diffusion scheme with b-values of 300, 650, 1000, and 2000 s/mm², corresponding to 4, 17, 39, and 68 diffusion-encoding directions, respectively. One non-diffusion-weighted (b = 0) volume was also included. The acquisition was performed using a single-shot echo-planar imaging (EPI) sequence with anterior-to-posterior (AP) phase encoding. Each diffusion volume consisted of 60 axial slices acquired with a 2 mm isotropic resolution (slice thickness = 2 mm, in-plane resolution = 2 × 2 mm), and the field of view (ReadoutFOV) was 220 mm. The total scan duration was approximately 6.5 minutes. Acquisition parameters for the diffusion imaging included TR = 2750 ms and TE = 79 ms.

Following data acquisition, the diffusion data were reconstructed in the MNI space using q-space diffeomorphic reconstruction (QSDR)^79^ to obtain the spin distribution function (SDF)^80^ in DSI Studio (Version Hou, 2024). A diffusion sampling length ratio of 1.25 was applied, with the output resolution in diffeomorphic reconstruction set to 2 mm isotropic. The tensor metrics were then calculated.

For fiber tractography, a deterministic fiber tracking algorithm^81^ was used, incorporating augmented tracking strategies^82^ to improve reproducibility. The quantitative anisotropy (QA) threshold was set to 0.12, and the angular threshold was set to 60 degrees. The step size was 1.00 mm, and tracks shorter than 10 mm or longer than 400 mm were discarded. A total of 5 million tracts were calculated for each subject. Shape analysis was conducted to derive shape metrics for the tractography.^82^

For full reproducibility, the parameter ID used in DSI Studio to configure these settings is 8FC2F53D9A99193Fba3Fb803Fcb2041bC843404B4Cca01cbaCDCC4C3Ec. This ID allows others to load the exact settings and parameters used in our analysis, ensuring that the tractography and other metrics can be reproduced using the same configurations.

The functional imaging was performed using a T2-weighted gradient echo-planar imaging (EPI)* sequence during the behavioral tasks. Four functional runs were acquired with the following parameters: TR = 535 ms, TE = 30 ms, flip angle = 46°, and voxel size = 3.8 mm × 3.8 mm × 4 mm. The field of view was 240 mm, and 32 slices were collected in an interleaved order with a slice thickness of 4 mm. A total of 3440 brain images were acquired during task performance.

#### 2.2.2 fMRI Pre-Processing Pipeline

The preprocessing pipeline for functional MRI data combined with tools from AFNI ^83, 84^ and FSL^85–87^ to ensure high-quality data for subsequent analysis^88, 89, 90^. The process began with denoising the fMRI data to reduce noise from physiological artifacts such as head motion and scanner drift, using AFNI’s dwidenoise command. This step resulted in denoised datasets for each run of the fMRI data. Following this, motion correction was performed using AFNI’s 3dvolreg, which registers each volume of the fMRI data to a reference volume within the session. After motion correction, the data were aligned to the MNI space using FSL’s FLIRT tool, ensuring that all data were in a common standard space for group-level comparisons.

To remove any motion-related artifacts, outlier detection was carried out using AFNI’s 3dToutcount, which computes the fraction of outlier voxels in each volume. A censoring procedure was then applied, excluding volumes where the fraction of outlier voxels exceeded a predefined threshold (0.1). Despiking was performed with AFNI’s 3dDespike, which removed brief, spurious signal fluctuations, or “spikes,” from the data.

Following despiking, slice timing correction was applied using 3dTshift to ensure temporal alignment across slices in each volume.

Time series were then extracted from predefined brain regions based on the AAL3 parcellation using AFNI’s 3dROIstats, which computes the average signal within each region of interest (ROI). These time series were saved as text files for further analysis. Signal-to-noise ratio (SNR) for each run was computed using AFNI’s 3dTstat to calculate the mean signal and 3dTproject to compute the standard deviation of the noise.

Additionally, global correlation averages (GCOR) were computed to assess overall data quality.

The degree of spatial blurring in the data was estimated using FSL’s 3dFWHMx, which calculates the full width at half maximum (FWHM) of the data’s spatial blurring. Following this, an extents mask was created using AFNI’s 3dmask_tool to identify valid brain regions with usable data across all volumes. The data were then registered to the MNI template (ENIGMA Template) using AFNI’s @auto_tlrc, which applies a transformation matrix to warp each subject’s data into standard space for group-level analysis.

To further improve data quality, Principal Component Analysis (PCA) was applied using AFNI’s 3dpc to remove non-neuronal signals, such as global signal fluctuations and motion-related noise. PCA regressors were generated from ventricular and brain regions and were used in subsequent regression analysis to remove unwanted variance from the data. Finally, the processed datasets were reviewed for quality control, and any remaining temporary files were removed to prepare the data for further analysis.

After the AFNI and FSL preprocessing steps, additional processing was performed in MATLAB to further refine the data. This included outlier correction, where extreme values in the time series that exceeded 5 standard deviations were identified and corrected. Detrending was applied to remove any linear trends from the data using MATLAB’s detrend function, ensuring that any slow drifts in the signal did not affect subsequent analyses. The time series data were then parsed by behavioral condition and time block, creating condition-specific time series data for each subject. This allowed for more detailed analysis of brain activity in relation to specific experimental conditions. The preprocessed time series data for each region of interest (ROI) in the AAL3 atlas were stored in structured files and saved for subsequent analysis.

### 2.3 Atlas Selection & AAL3 Parcellation

For the functional and structural connectivity analysis, the Automated Anatomical Labeling 3 (AAL3) atlas was selected due to its widespread use, and strong effect sizes in capturing brain structure-function relationships, especially when compared to other well-known atlases.^91^ The AAL3 atlas includes 166 parcellations, with critical task-relevant regions such as the orbitofrontal cortex, cerebellum, and thalamic nuclei, which are particularly relevant for video game studies investigating neural processes underlying cognitive functions like visuomotor decision-making. This makes it highly suitable for whole-brain analysis in our study.^92^ For better visual clarity and interpretability, we organized the regions of the AAL3 atlas into clear subdivisions Orbitofrontal, Occipital, Limbic System, Frontal, Temporal, Thalamus, Parietal, Basal Ganglia, Cerebellum, and Brain Stem (see Figure 6a) based on their known anatomical locations, while preserving individual regions in our analysis as displayed in Supplementary Figure 1. As the brain slices progress in Figure 6b, the organization of these regions becomes more apparent, revealing how these anatomical structures are spatially arranged. This clear organizational structure aids in interpreting the results of our analysis.

**Figure 6.**
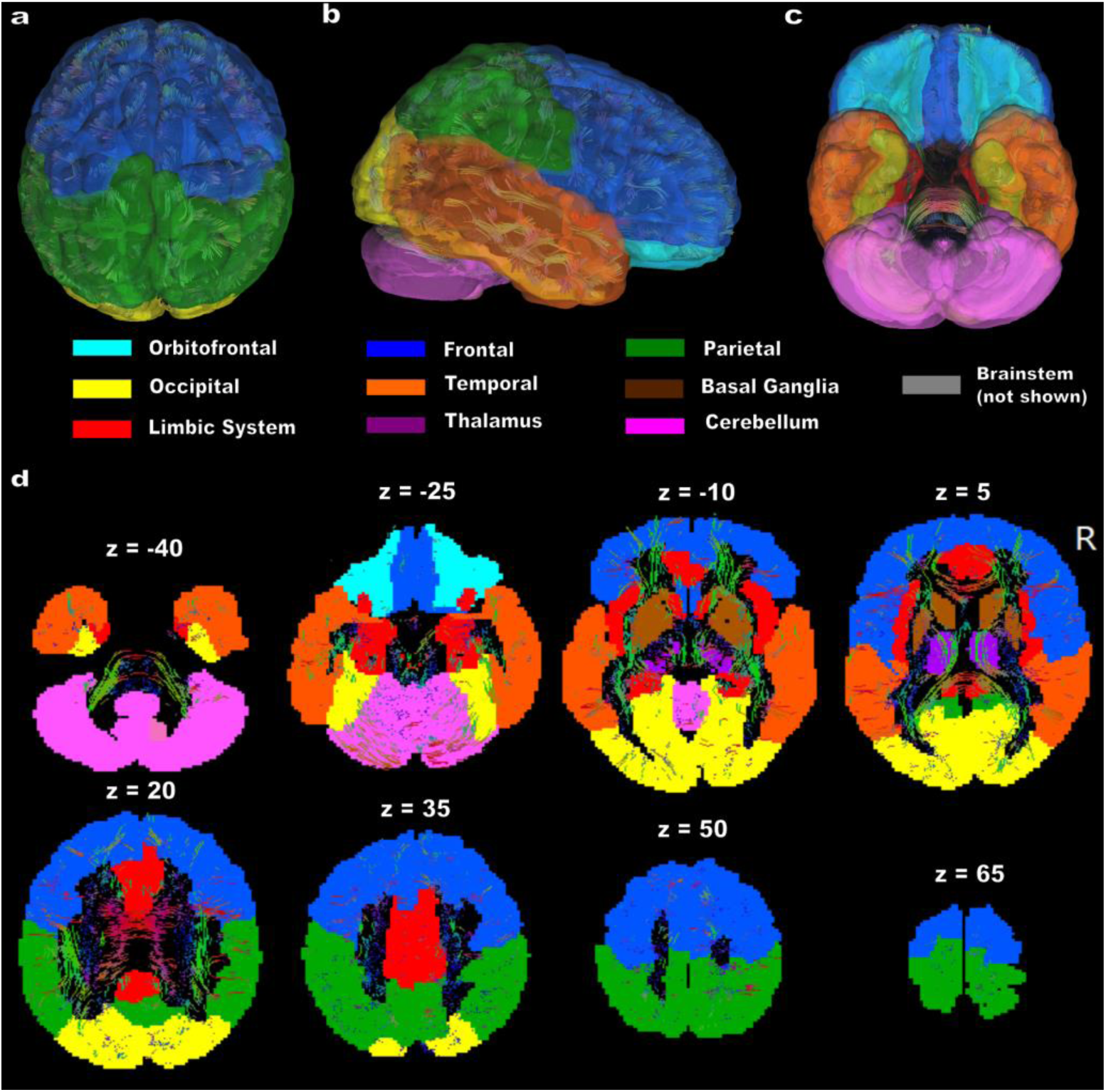
AAL3 Atlas Parcellation Categories for Connectivity Analysis. Visualization of the AAL3 atlas with anatomically grouped parcellations used in the connectivity analysis. **(a)** Superior view, **(b)** Right lateral view, **(c)** Inferior view. **(d)** Axial slices illustrate the parcellation structure along the z-axis. Colors correspond to distinct anatomical groups. The brainstem (gray) is not shown but is included in the analysis.

### 2.4 Connectivity and Graph Theoretic Analysis (SC-FC)

#### 2.4.1 Computation of SC-FC Connectivity

Structural-functional connectivity (SC-FC) was computed by combining structural and functional connectivity matrices. Functional connectivity (FC) was calculated using Pearson’s correlation coefficient between the time series of different brain regions, obtained from the AAL3 parcellation. The functional connectivity matrix was created using time series data extracted from predefined brain regions across all participants.

For the structural connectivity (SC) analysis, diffusion-weighted imaging (DWI) data were processed to derive a structural connectivity matrix based on deterministic tractography. This process involved the use of q-space diffeomorphic reconstruction (QSDR) and a deterministic fiber tracking algorithm to estimate the structural connectivity between brain regions. The result was a matrix that quantified the number and strength of the structural connections between each pair of regions in the brain To constrain the functional connectivity data by structural connectivity, a QA threshold of 0.12 was applied to the structural connectivity matrix (using the same threshold as the original tractography analysis). This threshold was used to binarize the QA connectivity matrix, effectively excluding weak or insignificant connections. Once this binarized mask was created for each participant, it was applied to the functional connectivity matrix, filtering the FC data by the structural connectivity constraints. This approach ensured that only the functional connections between regions with significant structural connectivity were retained for further analysis.

Once the SC-FC data were obtained for each subject, Fisher’s Z transformation was applied to the Pearson’s correlation coefficients to ensure valid and unbiased statistical comparisons, as Pearson’s correlation is non-linear and not normally distributed. Mean differences between groups were then computed on the *Z*-transformed data. The Mann-Whitney U rank sum test (*p* < 0.05) was used to determine significance, as it is a non-parametric method that allows for confident identification of significant differences, regardless of the underlying data distribution, minimizing the risk of false positives while maintaining sensitivity to true effects.^93^ To ensure the test assumptions were met, we verified that the degree of skewness in our data was comparable between groups. After significance testing, our data was transformed back into correlation coefficients for interpretability.

#### 2.4.2 SC-FC Graph-Theoretic Analysis

For the undirected graph-theoretic network analysis, we applied a 95% threshold to binarize the SC-FC data, retaining the top 95% of the strongest connections after applying the structural connectivity (SC) constraints. This approach was chosen to capture as much of the structural network as possible while ensuring that only valid, non-spurious connections were included.

Binarizing the network simplifies the analysis by focusing on the presence or absence of connections rather than their strength. Many recent studies discard link weights, as binary networks are in most cases simpler to characterize and have a more easily defined null model for statistical comparison^94^ making them more reliable for exploratory data analysis. This approach was particularly suited for examining whole-brain networks, allowing us to explore brain-wide network dynamics across many regions.

We considered both global and local graph-theoretic measures in our analysis, which were calculated using the Brain Connectivity Toolbox (BCT). ^94^ Node degree and local efficiency were calculated for each AAL3 region in both gamers and non-gamers. These graph-theoretic measures were used to assess the regional properties of network nodes, with node degree providing insight into the number of connections each region has, and local efficiency quantifying how effectively information is exchanged within each region’s local network

### 2.5 Connectivity and Graph Theoretic Analysis (SC-dFC)

#### 2.5.1 Computation of SC-dFC Connectivity

Structural-directed functional connectivity (SC-dFC) was computed by combining structural and directed functional connectivity matrices. Directed functional connectivity (dFC) was calculated using time-domain Granger causality (GC) defined as follows:

The time interval-specific GC (*I*) from region 2 (source) to region 1 (target) in the frequency domain is defined as

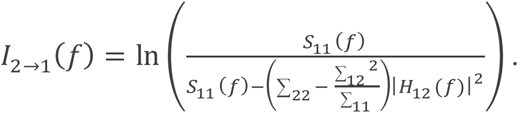

The frequency band-specific or time-domain equivalent GC (*F)* is then calculated using

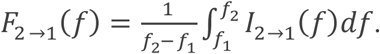

In the bivariate autoregressive model ^42, 95^. The noise covariance and transfer function matrices are denoted by “∑” and “H” respectively. The evaluation of the frequency band-specific or time-domain equivalent Granger causality (GC) was accessed in the range between *f*_1_= 0.05 Hz to *f*_2_ = 0.9 Hz, with a sampling rate of 1.87 Hz (*TR*^−1^).

The appropriate model order for the TGC analysis was determined by minimizing the spectral difference between the Granger-generated time series and the original signal, while maintaining sensitivity to trial-specific dynamics governed by the trial duration and the repetition time (TR). To preserve this sensitivity, the model order was constrained such that it did not exceed the number of time points within a trial. The maximum allowable model order, denoted mo_max_, is given by

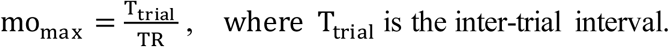

Under this constraint, a model order of 5 was selected, as it best minimized the spectral discrepancy in the whole-brain analysis while preserving sensitivity to task-related GC fluctuations. TGC matrices were computed between the AAL3 region time series using this model order, which served as the directed functional connectivity (dFC).

For structural connectivity (SC) analysis, diffusion-weighted imaging data were processed to derive a structural connectivity matrix using deterministic tractography. This involved q-space diffeomorphic reconstruction (QSDR) and a deterministic fiber tracking algorithm to estimate the number and strength of structural connections between each pair of brain regions.

To constrain the dFC data, a quantitative anisotropy (QA) threshold of 0.12 was applied to the SC matrix, consistent with the threshold used in the original tractography analysis. This threshold was used to binarize the QA-based connectivity matrix, excluding weak or insignificant connections. The resulting binary mask was then applied to each participant’s dFC matrix, allowing only directed functional connections between structurally connected regions to be retained for further analysis.

Importantly, applying the SC mask to the dFC data yields a measure of effective connectivity, which captures the directional and causal influence one brain region exerts on another, constrained by the underlying anatomical substrate^43^. Group differences were assessed using the Mann–Whitney U rank-sum test (*p* < 0.05).

#### 2.5.2 SC-dFC Graph-Theoretic Analysis

FC is inherently symmetric, meaning that a valid connection between two nodes (A, B) implies that the same connection, with the same magnitude, exists for (B, A). In contrast, dFC is inherently asymmetric, meaning that a connection from source A to target B does not imply the same or a similar connection from B to A. This asymmetry results in dFC networks exhibiting greater variability and a sparser structure compared to FC networks.

To address this sparsity, we applied a 10% threshold to dFC, retaining only the top 10% of the strongest and most reliable directed interactions. This threshold minimized the influence of invalid or spurious connections, while preserving as many effective connections as possible. By doing so, we ensured that the global density of the SC-dFC network remained comparable to that of the SC-FC network within the same subject. This thresholding strategy aligned with our goal of maintaining as many meaningful connections as possible to accurately characterize the SC-dFC network while excluding invalid and spurious connections.

We considered both global and local graph-theoretic measures in our analysis, which were also calculated using the Brain Connectivity Toolbox (BCT).^94^ Node degree and local efficiency were calculated for each AAL3 region in both gamers and non-gamers.

### 2.6 Assessing Behavioral Relevance

To assess the behavioral relevance of brain connectivity measures and functional network properties, we used Spearman correlation to examine the association between each brain connectivity measure (SC-FC or SC-dFC) and response time (RT) in the behavioral task. The correlations were calculated for both group-level comparisons and within-group analyses.

For group comparisons, we tested for significant differences using the Mann-Whitney U rank-sum test with significance defined as *p* < 0.05 and |*r*| ≥ 0.2. If a significant difference was observed, we plotted the corresponding brain-behavior correlation graph for the entire group, as well as separate graphs for each group (gamers and non-gamers) to explore within-group relationships. A positive correlation is associated with a brain measure that tracks slower response times, while a negative correlation is associated with faster response times.

This approach allowed us to identify brain regions and connectivity patterns that were significantly associated with response time differences across groups, as well as the extent to which these relationships persisted within each group.

## SUPPLEMENTARY FIGURES

**SUPPLEMENTARY FIGURE 1.**
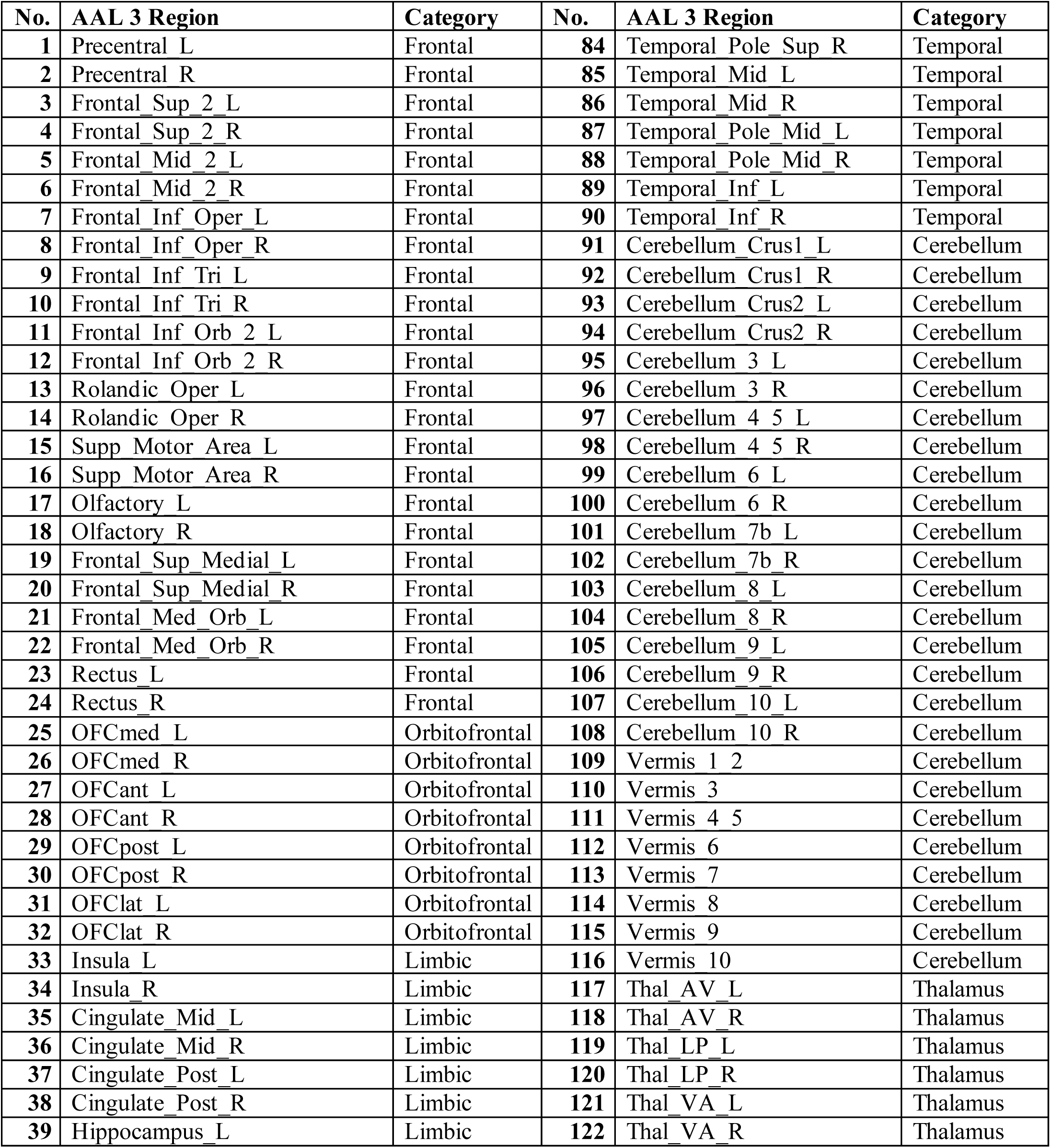
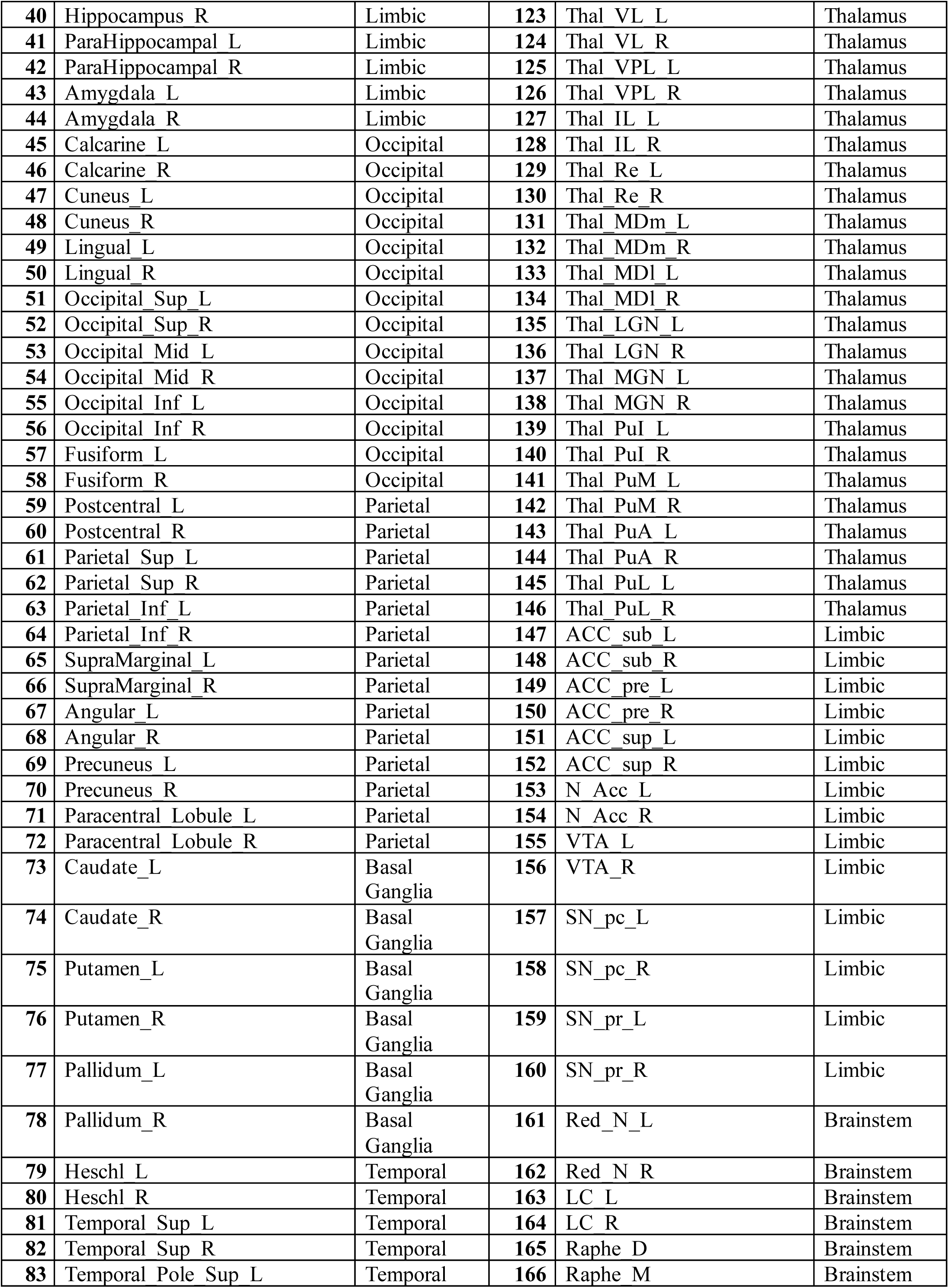
AAL 3 Region Parcellation Categories by Major Anatomical Groups. *Original numbers in AAL2 for the anterior cingulate cortex (ACC) and thalamus are left empty in AAL3, as those voxels were substituted by the new subdivisions for the Thalamic nuclei and ACC.*^92^ *These regions are removed here and the index has been adjusted accordingly to the correct 166 total parcellations categorized by major anatomical groups*.

## Author Contributions

**Kyle Cahill:** Conceptualization, Methodology, Software, Formal analysis, Writing – original draft, review & editing. **Mukesh Dhamala:** Conceptualization, Methodology, Software, Supervision, Funding acquisition, Writing – review & editing.

## Data Availability Statement

All data that support the findings of the study as well as the custom analysis scripts can be found in OSF (*link will be public after publication*)

## Ethics Statement

This research was conducted in accordance with all relevant ethical guidelines, including informed consent procedures, participant confidentiality, and adherence to the principles outlined by the Institutional Review Board at Georgia State University. All participants provided written informed consent prior to data collection, and their identities were protected throughout the study.

## Acknowledgments

This work was funded by two internal grant awards of Brains and Behavior Program & the Center for Advanced Brain Imaging to M.D. Bhim Adhikari for providing fMRI preprocessing pipeline. Tim Jordan for initial data collection.

